# Disrupting HIV-1 capsid formation causes cGAS sensing of viral DNA

**DOI:** 10.1101/838011

**Authors:** Rebecca P. Sumner, Lauren Harrison, Emma Touizer, Thomas P. Peacock, Matthew Spencer, Lorena Zuliani-Alvarez, Greg J. Towers

## Abstract

Detection of viral DNA by cyclic GMP-AMP synthase (cGAS) is a first line of defence leading to the production of type-I interferon (IFN). As HIV-1 is not a strong inducer of IFN we have hypothesised that its capsid cloaks viral DNA from cGAS. To test this we generated defective viral particles by treatment with HIV-1 protease inhibitors or by genetic manipulation of *gag*. These viruses had defective Gag cleavage, reduced infectivity and diminished capacity to saturate TRIM5α. Importantly, unlike wild-type HIV-1, infection with cleavage defective HIV-1 triggered an IFN response in THP-1 cells and primary human macrophages that was dependent on viral DNA and cGAS. Infection in the presence of the capsid destabilising small molecule PF-74 also induced a cGAS-dependent IFN response. These data demonstrate a protective role for capsid and suggest that antiviral activity of capsid- and protease-targeting antivirals may benefit from enhanced innate and adaptive immunity *in vivo*.

## Introduction

The innate immune system provides the first line of defence against invading pathogens such as viruses. Cells are armed with pattern recognition receptors (PRRs) that recognise pathogen-associated molecular patterns (PAMPs), such as viral nucleic acids, and lead to the activation of a potent antiviral response in the form of secreted interferons (IFNs), proinflammatory cytokines and chemokines, the expression of which is driven by the activation of key transcription factors such as IFN regulatory factor 3 (IRF3) and nuclear factor kappa-light-chain-enhancer of activated B cells (NF-κB) (Chow, Franz et al., 2015). For HIV-1 a number of cytosolic PRRs have been demonstrated to contribute to the detection of the virus in infected cells including DNA sensors cyclic GMP-AMP synthase (cGAS) (Gao, Wu et al., 2013, Lahaye, Satoh et al., 2013, Rasaiyaah, Tan et al., 2013), IFI16 (Jakobsen, Bak et al., 2013, Jonsson, Laustsen et al., 2017) and PQBP1 (Yoh, Schneider et al., 2015) and RNA sensors DDX3 (Gringhuis, Hertoghs et al., 2017) and also MDA5, although only in the circumstance where the genome lacked 2′-*O*-methylation by 2′-*O*-methyltransferase FTSJ3 (Ringeard, Marchand et al., 2019). The best studied of these sensors is cGAS, which upon binding double-stranded DNA, such as HIV-1 reverse transcription (RT) products, produces second messenger 2’3’-cGAMP (Ablasser, Goldeck et al., 2013, Sun, Wu et al., 2013, Wu, Sun et al., 2013) that binds and induces phosphorylation of ER-resident adaptor protein STING and its translocation to perinuclear regions (Tanaka & Chen, 2012). Phosphorylation of STING provides a platform for the recruitment of TBK1 and IRF3 leading to IRF3 phosphorylation and its subsequent translocation to the nucleus to drive expression of IFN and IFN stimulated genes (ISGs) (Liu, Cai et al., 2015). Activation of STING by 2’3’-cGAMP also activates IKK and the transcription of NF-κB-dependent genes (Ishikawa & Barber, 2008).

Of course, detection of infection by sensing is not universal and viruses are expected to hide their PAMPs and typically have mechanisms to antagonise specific sensors and downstream restriction factors. Work from our lab (Rasaiyaah et al., 2013) and others (Cingoz & Goff, 2019) has demonstrated that primary monocyte-derived macrophages (MDMs) can be infected by wild-type (WT) HIV-1 without significant innate immune induction. However, MDM sense HIV-1 if, for example, mutations are made in the viral capsid to prevent the recruitment of cellular cofactors such as CPSF6 and cyclophilin A (Rasaiyaah et al., 2013) or after depletion of the cellular exonuclease TREX1 (Rasaiyaah et al., 2013, Yan, Regalado-Magdos et al., 2010). This sensing was found to be dependent on viral reverse transcription (RT) and the cellular DNA sensing machinery cGAS and STING. In addition to recruitment of cofactors, a variety of evidence suggests that capsid remains intact in the cytoplasm to protect the process of viral DNA synthesis, preventing degradation of RT products by cellular nucleases such as TREX1 and from detection by DNA sensors (Burdick, Delviks-Frankenberry et al., 2017, Francis & Melikyan, 2018).

Here we have tested the hypothesis that an intact capsid is crucial for innate immune evasion by disrupting the process of viral particle maturation, either biochemically using protease inhibitors (PIs), or genetically, by mutating the cleavage site between the capsid protein and spacer peptide 1. The resulting viral particles had defective Gag cleavage, reduced infectivity and, unlike wild-type HIV-1, activated an IFN-dependent innate immune response in THP-1 cells and primary human macrophages. This innate response was mostly dependent on viral DNA synthesis and the cellular sensors cGAS and STING. Defective viruses were less able to saturate restriction by TRIM5α indicating a reduced ability to bind this restriction factor, likely due to aberrant particle formation. Finally, we show that the capsid binding small molecule inhibitor PF-74, which has been proposed to accelerate capsid opening (Marquez, Lau et al., 2018), also induces HIV-1 to activate an innate response in THP-1 cells, that is dependent on cGAS. Together these data support the hypothesis that the viral capsid plays a physical role in protecting viral DNA from the cGAS/STING sensing machinery in macrophages and that disruption of Gag cleavage and particle maturation leads to aberrant capsid formation and activation of an IFN response that may be harnessed therapeutically *in vivo* during PI treatment of HIV-1.

## Results

### Protease inhibitor treatment of HIV-1 leads to innate immune induction in macrophages

To test the hypothesis that intact viral capsids protect HIV-1 DNA from detection by DNA sensors we sought to generate defective viral particles by disrupting capsid maturation. The protease inhibitor (PI) class of anti-retrovirals block the enzymatic activity of the viral protease, preventing Gag cleavage and proper particle formation, as observed by electron microscopy (Muller, Anders et al., 2009, Schatzl, Gelderblom et al., 1991). By producing VSV-G-pseudotyped HIV-1ΔEnv.GFP (LAI strain (Peden, Emerman et al., 1991) with the Nef coding region replaced by GFP, herein called HIV-1 GFP) in the presence of increasing doses of the PI lopinavir (LPV, up to 100 nM) we were able to generate viral particles with partially defective Gag cleavage, as assessed by immunoblotting of extracted viral particles detecting HIV-1 CA protein (Fig. 1A). At the highest dose of LPV (100 nM) increased amounts of intermediate cleavage products corresponding to capsid and spacer peptide 1 (CA-SP1), matrix and CA (MA-CA), MA, CA, SP1 and nucleocapsid (MA-NC) were particularly evident along with increased amounts of full length uncleaved Gag (Fig. 1A, Suppl. Fig. 2A). Uncleaved CA-SP1 was also evident at 30 nM LPV. As expected, defects in Gag cleavage were accompanied by a reduction in HIV-1 GFP infectivity in both phorbol myristyl acetate (PMA)-treated THP-1 (Fig. 1B) and U87 cells (Fig. 1C). For the highest dose of LPV this corresponded to a 24- and 48-fold defect in infectivity in each cell type respectively. Viral titres were calculated according to the number of genomes, assessed by qPCR (see Methods), to account for small differences in viral production between conditions. These differences were no more than 2-fold from untreated virus.

**Fig. 1.**
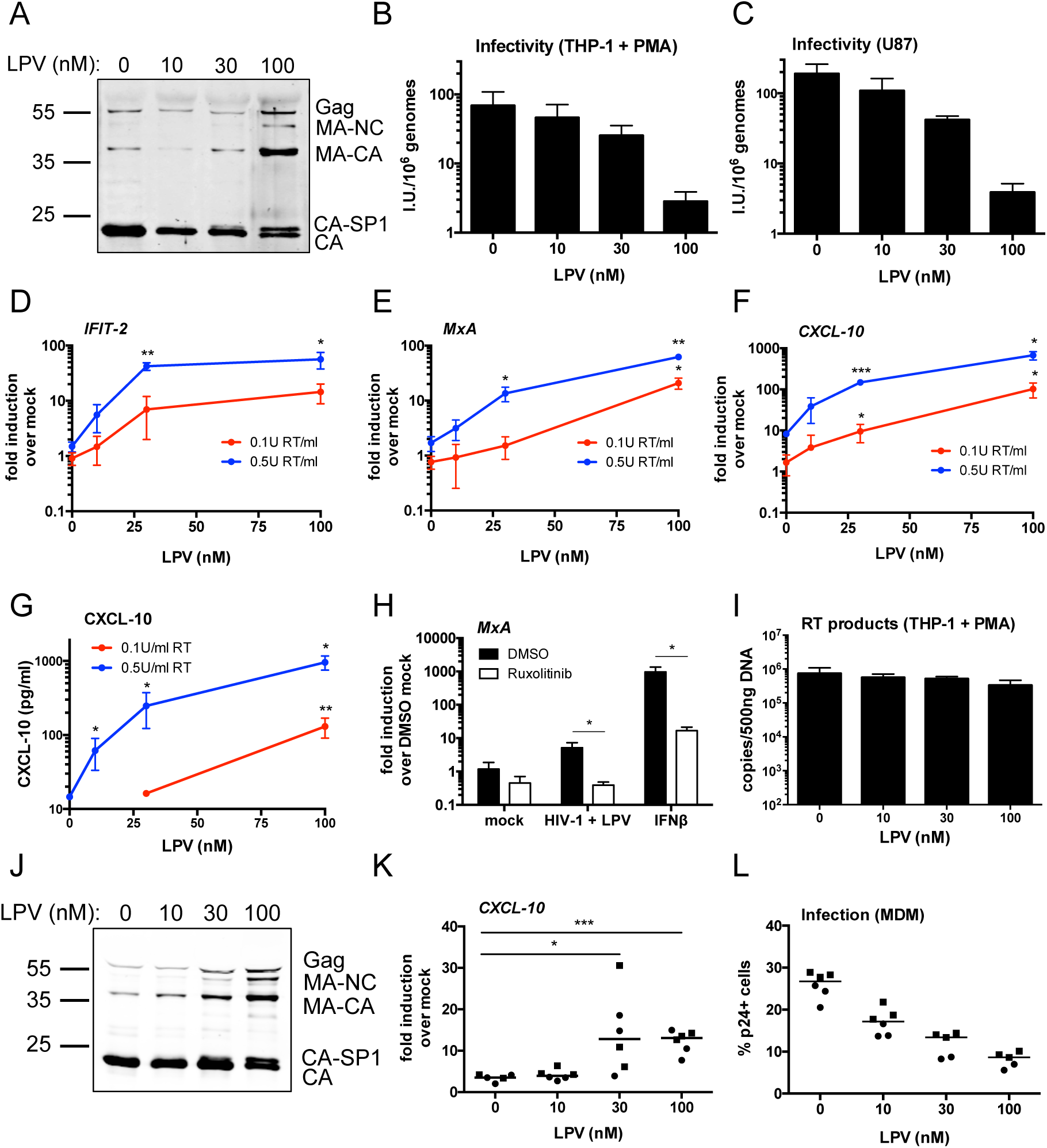
PI treatment induces HIV-1 to trigger an ISG response in macrophages. (A) Immunoblot of HIV-1 GFP virus particles (2×10^11^ genomes) produced in HEK293T cells in the presence of increasing doses of protease inhibitor lopinavir (LPV, 0-100 nM) and then purified through a 20 % sucrose cushion. The blot was probed with an anti-HIV-1 p24 antibody. Molecular mass markers (in kDa) are indicated on the left and Gag cleavage products on the right. (B-C) Titration of LPV-treated HIV-1 GFP viruses on PMA-treated (50 ng/ml, 48 h) THP-1 shSAMHD1 (B) or U87 (C) cells. Infectious units per ml (IU/ml) were calculated by enumeration of GFP-positive cells by flow cytometry at 48 h post-transduction. These data were then normalised to genomes/ml data calculated by qPCR to account for variations in viral production. Data are presented as mean ± SD from three separate titrations performed in singlet. (D-F) ISG qPCR from PMA-treated (50 ng/ml, 48 h) THP-1 shSAMHD1 cells that had been transduced for 24 h with LPV-treated HIV-1 GFP viruses (0.1 U RT/ml red line, 0.5 U RT/ml blue line). Expression of *IFIT-2* (D), *MxA* (E) and *CXCL-10* (F) was normalised to an internal control (*GAPDH*) and these values were then normalised to those for the non-transduced mock cells, yielding the fold induction over mock. (G) Level of CXCL-10 protein in the cell supernatants from D-F was measured by ELISA. Data for D-G are presented as mean ± SD of triplicate data repeated at least three times. (H) ISG qPCR from PMA-treated (50 ng/ml, 48 h) THP-1 shSAMHD1 cells that had been transduced for 24 h with 0.2 U RT/ml HIV-1 GFP that had been treated with 30 nM LPV, or stimulated with 1 ng/ml IFNβ as a control, in the presence or absence (DMSO control) of 2 µM ruxolitinib. Expression of *MxA* was normalised to an internal control (*GAPDH*) and these values were then normalised to those for the non-transduced, DMSO-treated mock cells, yielding the fold induction over DMSO mock. Data are presented as mean ± SD of triplicate data repeated at least twice. (I) RT products from PMA-treated (50 ng/ml, 48 h) THP-1 shSAMHD1 cells that had been transduced for 24 h with 0.3 U RT/ml LPV-treated HIV-1 GFP viruses. Cells were lysed for DNA extraction and then used for qPCR analysis using primers for GFP. Data are presented as mean ± SD of triplicate data repeated at least twice. (J) Immunoblot of R9 BaL virus particles (2×10^11^ genomes) that were produced in HEK293T cells in the presence of increasing doses of LPV (0-100 nM) and then purified through a 20 % sucrose cushion. The blot was probed with an anti-HIV-1 p24 antibody. Molecular mass markers (in kDa) are indicated on the left and Gag cleavage products on the right. (K) ISG qPCR from primary monocyte-derived macrophages (MDM) that had been infected for 24 h with LPV-treated R9 BaL viruses (0.2 U RT/ml). Expression of *CXCL-10* (K) was normalised to an internal control (*GAPDH*) and these values were then normalised to those for the non-infected mock cells, yielding the fold induction over mock. (L) infection levels of cells from (K). Cells were stained with a FITC-labelled anti-p24 antibody and analysed by flow cytometry. Data presented for (K) and (L) are collated from two donors (represented by circles and squares), performed in triplicate. Horizontal line represents the median. Statistical analyses were performed using the Student’s t-test, with Welch’s correction where appropriate and comparing to the 0 nM LPV condition. * *P*<0.05, ** *P*<0.01, *** *P*<0.001. See also Suppl. Fig. 2 and 3.

To test the visibility of PI inhibited viruses to innate sensing responses we generated a THP-1 cell line that was stably depleted for the HIV restriction factor SAMHD1 (Suppl. Fig. 1A). Monocytic THP-1 cells can be differentiated into macrophage-like adherent cells by treatment with PMA, yielding a cell line that is highly competent for innate immune sensing, including DNA sensing. Differentiation of THP-1 normally leads to SAMHD1 activation by dephosphorylation and potent restriction of HIV-1 infection (Cribier, Descours et al., 2013). SAMHD1 depletion effectively relieved this restriction and allowed HIV-1 GFP infection (Suppl. Fig. 1A, B). SAMHD1-depleted THP-1 cells (herein referred to as THP-1 shSAMHD1 cells) remained fully competent for innate immune sensing and produced interferon-stimulated genes (ISGs) and inflammatory chemokines including *CXCL-10*, *IFIT-2* (also known as *ISG54*) and *CXCL-2* in response to a range of stimuli, including transfection of herring-testis DNA (HT-DNA), exposure to 2’3’-cGAMP and infection by Sendai virus (Suppl. Fig. 1C-E).

Infection of PMA-treated THP-1 shSAMHD1 cells with HIV-1 GFP that had been produced in the presence of increasing doses of LPV led to a virus and LPV dose-dependent increase in the expression of ISGs *CXCL-10*, *IFIT-2* and *MxA* at the mRNA level (Fig. 1D-F), and CXCL-10 protein secretion (Fig. 1G). In agreement with previous reports in primary macrophages (Cingoz & Goff, 2019), HIV-1 GFP produced in the absence of LPV induced very little, or no ISG expression in THP-1 cells at the doses tested, consistent with the hypothesis that HIV-1 shields its PAMPs from cellular PRRs (see Fig 1D-G, 0 nM drug dose). Virus dose in these experiments was normalised according to RT activity, as measured by SG-PERT (see Methods), which differed no more than 5-fold in the LPV-treated versus untreated virus. Infection levels in differentiated THP-1 cells were approximately equivalent between the various LPV doses tested (Suppl. Fig. 2B) because HIV-1 GFP infection of THP-1 is maximal at about 70 % GFP positivity (Pizzato, McCauley et al., 2015). Similar results were obtained with the PI darunavir (DRV); treatment of HIV-1 GFP with increasing doses of DRV (up to 50 nM) led to defects in Gag cleavage (Suppl. Fig. 3A), decreased infectivity (Suppl. Fig. 3B, F) and at 12.5 and 25 nM DRV activated an ISG response in PMA-treated THP-1 shSAMHD1 cells (Suppl. Fig. 3D-E).

To test whether LPV-treated HIV-1-induced ISG expression in THP-1 cells depended on IFN production or direct activation of ISGs, infections were repeated in the presence of the JAK1/2 inhibitor ruxolitinib (Quintas-Cardama, Vaddi et al., 2010). Activation of STAT transcription factors downstream of IFN receptor engagement requires phosphorylation by JAKs and hence ruxolitinib inhibits IFN signalling (Fig. 1H). Induction of *MxA* (Fig. 1H) and *CXCL-10* (Suppl. Fig. 2C) expression by LPV-treated HIV-1 GFP was severely reduced in the presence of ruxolitinib, indicating that induction of ISG expression in these experiments requires an infection-driven type I IFN response. Treatment of cells with type I IFN provided a positive control for ruxolitinib activity (Fig. 1H, Suppl. Fig. 2C). Importantly, viral DNA production, measured by qPCR in infected PMA-treated THP-1 shSAMHD1 cells, was not changed by increasing LPV dose suggesting that PI inhibited HIV-1 makes normal levels of DNA but fails to protect PAMPs from innate immune sensors (Fig. 1I).

To test whether PI inhibition of HIV-1 caused similar innate immune activation in primary human cell infection we turned to HIV-1 R9 (Ba-L Env) infection of primary human macrophages. Production of R9 (BaL-Env) in HEK293T cells in the presence of 10-100 nM LPV induced the expected defects in Gag cleavage (Fig. 1J) and infectivity (Suppl. Fig. 2D and E) as observed with VSV-G-pseudotyped HIV-1 GFP (Fig. 1A-C). Furthermore, virus produced in the presence of 30 and 100 nM LPV induced the expression of *CXCL-10* on infection of primary MDM, whereas virus grown in the absence of LPV, or at low LPV concentrations (10 nM), induced very little *CXCL10* expression (Fig. 1K). Increasing concentrations of LPV during HIV-1 production led to a decrease in MDM infection, read out by p24 positivity, in these experiments (Fig. 1L). Together, these data demonstrate that infection by PI-treated HIV-1 induces an IFN-dependent innate immune response in PMA-treated THP-1 cells and primary human MDM that is not observed on infection with untreated virus.

### HIV-1 bearing Gag cleavage mutations also induces innate immune activation

Producing virus in the presence of PI suppresses Gag cleavage at multiple sites. Previous work suggested that inhibition of the CA-SP1 cleavage site was particularly toxic to infectivity and particles were defective with irregular partial polyhedral structures (Mattei, Tan et al., 2018, Muller et al., 2009). Concordantly our data show a defect in cleavage at the CA-SP1 site in the presence of LPV (Fig. 1A, J) or DRV (Suppl. Fig. 3A). Importantly, the presence of even small proportions of CA-SP1 cleavage mutant exerted trans-dominant negative effects on HIV-1 particle maturation (Muller et al., 2009). To test whether a CA-SP1 cleavage defect can cause HIV-1 to trigger innate sensing we prepared chimeric VSV-G pseudotyped HIV-1 GFP viruses by transfecting 293T cells with varying ratios of WT HIV-1 GFP and HIV-1 GFP with CA-SP1 Gag mutant L363I M367I (Checkley, Luttge et al., 2010, Wiegers, Rutter et al., 1998). Increasing the proportion of the ΔCA-SP1 mutant increased the presence of uncleaved CA-SP1 detected by immunoblot (Fig. 2A). Defective cleavage was accompanied by a modest decrease in infectivity on U87 cells (Fig. 2B).

**Fig 2:**
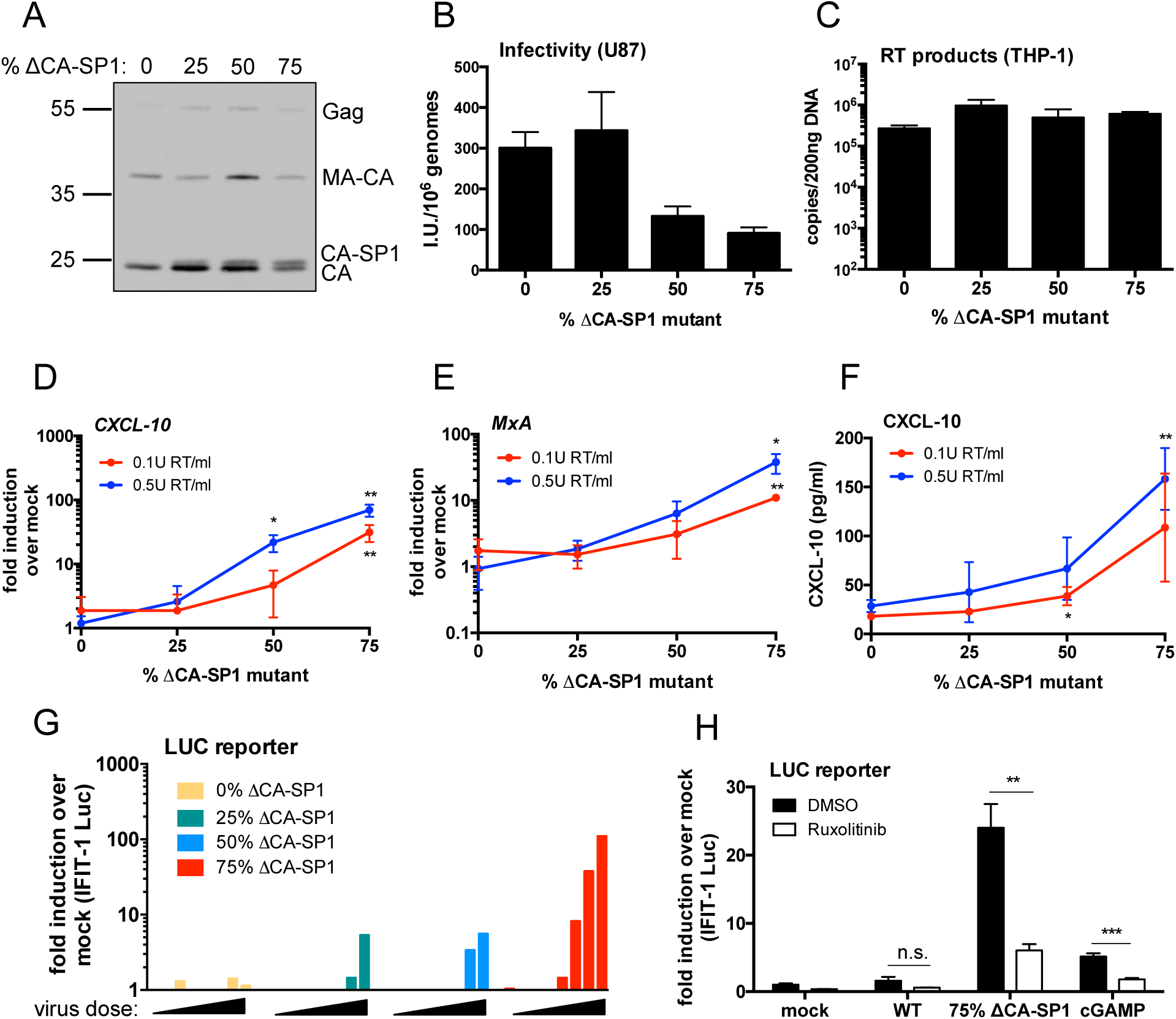
HIV-1 with Gag protease cleavage mutation induces ISGs in THP-1 cells. (A) Immunoblot of HIV-1 GFP virus particles (2×10^11^ genomes) produced in HEK293T cells with varying proportions of ΔCA-SP1 protease cleavage mutation and then purified through a 20 % sucrose cushion. The blot was probed with an anti-HIV-1 p24 antibody. Molecular mass markers (in kDa) are indicated on the left and Gag cleavage products on the right. (B) Titration of HIV-1 GFP ΔCA-SP1 viruses on U87 cells. Infectious units per ml (IU/ml) were calculated by enumeration of GFP-positive cells by flow cytometry. These data were then normalised to genomes/ml data calculated by qPCR to account for variations in viral production. Data are presented as mean ± SD from three separate titrations performed in singlet. (C) RT products from THP-1 cells that had been transduced for 24 h with 6×10^9^ genomes/ml (approx. 0.5 U RT/ml) HIV-1 GFP ΔCA-SP1 viruses. Cells were lysed for DNA extraction and then used for qPCR analysis using primers for GFP. Data are presented as mean ± SD of triplicate data repeated at least twice. (D-E) ISG qPCR from PMA-treated (50 ng/ml, 48 h) THP-1 shSAMHD1 cells that had been transduced for 24 h with HIV-1 GFP ΔCA-SP1 viruses (0.1 U RT/ml red line, 0.5 U RT/ml blue line). Expression of *CXCL-10* (D) and *MxA* (E) was normalised to an internal control (*GAPDH*) and these values were then normalised to those for the non-transduced mock cells, yielding the fold induction over mock. (F) Level of CXCL-10 protein in the cell supernatants from D-E was measured by ELISA. Data for D-F are presented as mean ± SD of triplicate data repeated at least three times. (G) IFIT-1 reporter activity from monocytic THP-1-IFIT-1 cells transduced for 24 h with varying doses of HIV-1 GFP ΔCA-SP1 viruses (0.016 – 0.2 U RT/ml). Gaussia luciferase activity in the supernatant was measured and normalised to mock transduced cells to generate a fold induction. Data are shown as individual measurements and are representative of at least two repeats. (H) IFIT-1 reporter activity from monocytic THP-1-IFIT-1 cells transduced with HIV-1 GFP containing either 0 % (WT) or 75 % ΔCA-SP1 mutant, or stimulated with 4 µg/ml cGAMP as a control, in the presence or absence (DMSO control) or 2 µM ruxolitinib. Gaussia luciferase activity in the supernatant was measured and normalised to DMSO-treated mock cells, yielding the fold induction over DMSO mock. Data are presented as mean ± SD of triplicate data repeated at least three times. Statistical analyses were performed using the Student’s t-test, with Welch’s correction where appropriate and comparing to the 0 % ΔCA-SP1 virus (D-F) or the DMSO control (H). * *P*<0.05, ** *P*<0.01, *** *P*<0.001. See also Suppl. Fig. 4.

As with HIV-1 GFP produced in the presence of PIs, infection of PMA-treated THP-1 shSAMHD1 cells with the HIV-1 GFP ΔCA-SP1 mutants led to a ΔCA-SP1 dose-dependent increase in the expression of *CXCL-10* (Fig. 2D) and *MxA* mRNA (Fig. 2E), and CXCL-10 at the protein level (Fig. 2F). Induction was not explained by differences in the amount of viral DNA in infected cells and similar levels of viral DNA (Fig. 2C) and infection (Suppl. Fig. 4A) were observed at the viral doses tested. Virus dose in these experiments was normalised according to RT activity, which differed no more than 5-fold between viruses. Cleavage defective viruses, and not wild type virus, also induced dose-dependent luciferase expression from an undifferentiated THP-1 cell line that had been modified to express Gaussia luciferase under the control of the *IFIT-1* (also known as *ISG56*) promoter, herein called IFIT1-luc (Mankan, Schmidt et al., 2014) (Fig. 2G, Suppl. Fig. 4B). IFIT1-luc is both IRF-3- and IFN-sensitive (Mankan et al., 2014). HIV-1 bearing ΔCA-SP1 mutant also induced a type I IFN response, evidenced by suppression of IFIT1-luc by ruxolitinib (Fig. 2H). In the IFIT1-luc cells ΔCA-SP1 mutation did not impact infection levels (Suppl. Fig. 4A-C) and neither did ruxolitinib treatment (Suppl. Fig. 4C). We propose that during single round infection the virus has already integrated by the time IFN is produced, thus explaining why ruxolitinib has no impact on the percentage of GFP positive cells. Together these data support our hypothesis that disruption of Gag maturation yields viral particles that fail to shield PAMP from innate sensors.

### Maximal innate immune activation by maturation defective viruses is dependent on viral DNA synthesis

To determine whether viral DNA synthesis is required for HIV-1 bearing ΔCA-SP1 to trigger sensing we infected THP-1 IFIT1-luc cells with HIV-1 75% ΔCA-SP1 in the presence of reverse transcriptase inhibitor neviripine and assessed sensing by measuring IFIT1-luc expression and CXCL10 secretion. As expected, infectivity was severely diminished by 5 µM neviripine (Suppl. Fig. 5A) and both luciferase (Fig. 3A) and CXCL-10 (Fig. 3B) secretion was completely inhibited suggesting that viral DNA synthesis is required to activate sensing. Concordantly, expression of ISGs *IFIT-2* (Fig. 3C, Suppl. Fig. 5B) and *MxA* (Fig. 3D, Suppl. Fig. 5B) induced by HIV-1 75% ΔCA-SP1 was also abolished in the presence of neviriprine. A small, but statistically significant reduction in luciferase (Fig. 3A) and CXCL-10 (Fig. 3B) secretion was observed in the presence of the integrase inhibitor raltegravir, although this was not observed in every experiment (Fig 3C-D). We conclude that viral DNA is the active PAMP and this notion was also supported by the observation that mutation D185E in the RT active site (HIV-1 ΔCA-SP1 RT D185E) also reduced activation of IFIT-1 luc expression (Fig. 3E) and CXCL10 secretion (Fig. 3F) on infection of the THP-1 IFIT-1 reporter cells. Mutation D116N of the viral integrase (HIV-1 ΔCA-SP1 INT D116N) impacted neither luciferase induction (Fig. 3E) or CXCL-10 (Fig. 3F) secretion.

**Fig. 3.**
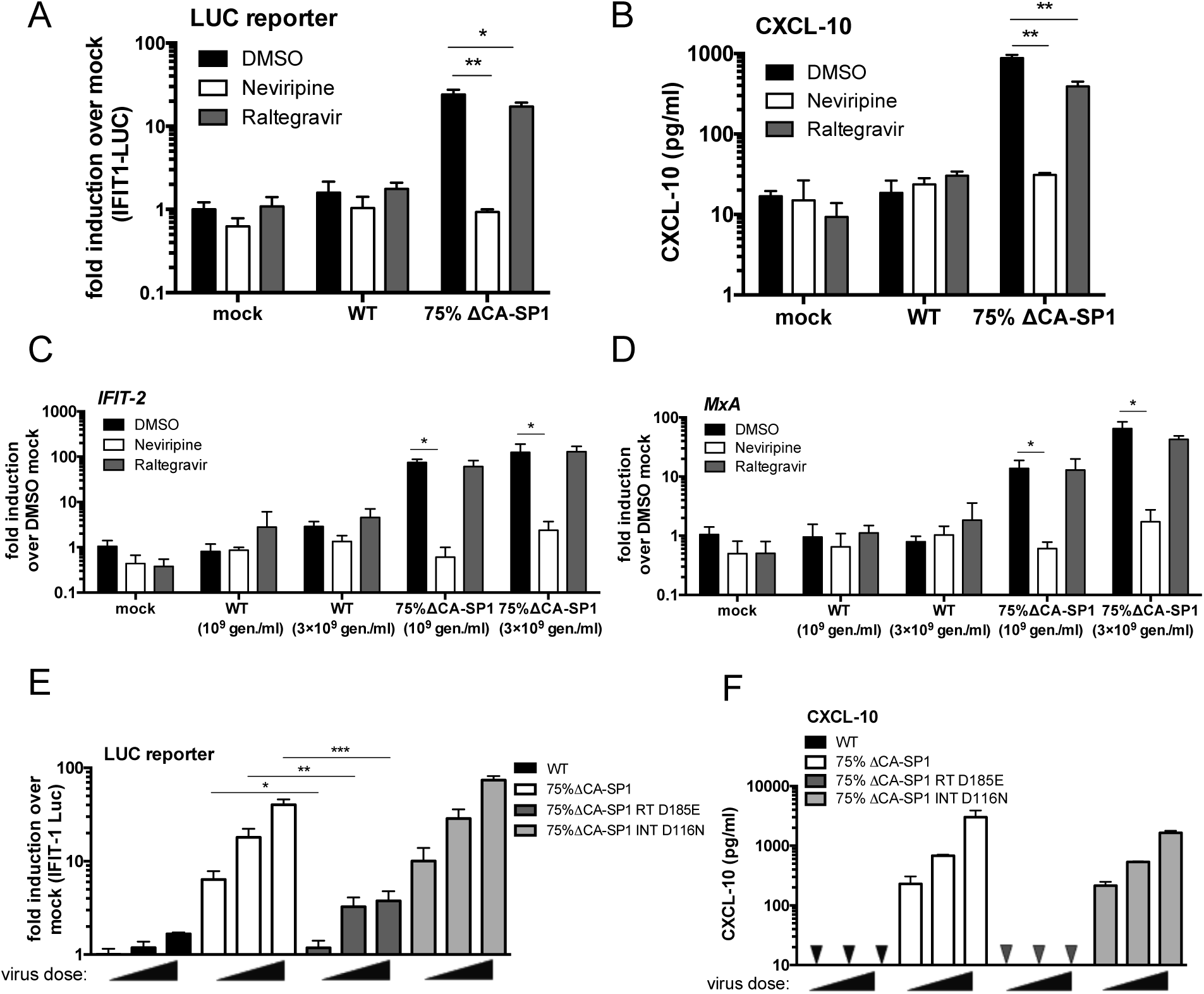
Innate immune activation is RT-dependent. (A) IFIT-1 reporter activity from THP-1-IFIT-1 cells transduced for 24 h with HIV-1 GFP containing 0 % (WT) or 75 % ΔCA-SP1 mutant (1 U RT/ml) in the presence or absence (DMSO control) of 5 µM neviripine or 10 µM raltegravir. Gaussia luciferase activity in the supernatant was measured and normalised to mock transduced cells to generate a fold induction. (B) Level of CXCL-10 protein in the cell supernatants from (A) was measured by ELISA. (C-D) ISG qPCR from THP-1-IFIT-1 cells that had been transduced for 24 h with increasing doses of 0 % (WT) or 75 % ΔCA-SP1 mutant (10^9^ ^and^ 3×10^9^ genomes/ml) in the presence or absence (DMSO control) of 5 µM neviripine or 10 µM raltegravir. Expression of *IFIT-2* (C) and *MxA* (D) was normalised to an internal control (*GAPDH*) and these values were then normalised to those for the non-transduced mock cells, yielding the fold induction over mock. (E) IFIT-1 reporter activity from THP-1-IFIT-1 cells transduced for 24 h with increasing doses of HIV-1 GFP containing 0 % ΔCA-SP1 (WT), 75 % ΔCA-SP1, 75 % ΔCA-SP1 carrying a mutation in reverse transcriptase (75 % ΔCA-SP1 RT D185E) or 75 % ΔCA-SP1 carrying a mutation in integrase (75 % ΔCA-SP1 INT D116N) (3.75×10^9^, 7.5×10^9^ and 1.5×10^10^ genomes/ml). (F) Level of CXCL-10 protein in the cell supernatants from B was measured by ELISA. Triangles indicate CXCL-10 not detected. Data are presented as mean ± SD of triplicate data repeated 2-3 times. Statistical analyses were performed using the Student’s t-test, with Welch’s correction where appropriate. * *P*<0.05, ** *P*<0.01, *** *P*<0.001. See also Suppl. Fig. 5.

Surprisingly neither treatment with 10 µM raltegravir (Suppl. Fig. 5A, B) or infection with HIV-1 ΔCA-SP1 INT D116N (Suppl. 5C) led to a reduction in GFP positivity in monocytic THP-1 cells. Importantly GFP expression was lost in parallel infection of PMA-treated THP-1 cells (Suppl. Fig. 5D, E) confirming that integration was indeed suppressed by 10 µM raltegravir or D116N integrase mutation. We propose that the GFP positivity observed in monocytic THP-1 cells in the presence of raltegravir, or by ΔCA-SP1 INT D116N, is due to expression from 2’-LTR circles that has been observed in other cell types (Bonczkowski, De Scheerder et al., 2016, Van Loock, Hombrouck et al., 2013).

### Viral DNA of maturation defective HIV-1 is sensed by cGAS and STING

To investigate which innate sensors were involved in detecting cleavage defective HIV-1, we infected cells that had been genetically manipulated by CRISPR/Cas 9 technology to lack the DNA sensing component proteins cGAS (Invivogen) or STING (Tie, Fernandes et al., 2018), or the RNA sensing component MAVS (Tie et al., 2018). As expected, STING-/- cells did not respond to transfected Herring Testis (HT)-DNA but ISG induction was maintained in response to the RNA mimic poly I:C (Fig. 4A). MAVS-/- cells showed the opposite phenotype, responding to poly I:C, but not HT-DNA (Fig. 4A). As expected, Dual IRF reporter THP-1 cells, knocked out for cGAS (Invivogen), responded normally to poly I:C, LPS and cGAMP but not transfected HT-DNA (Fig. 4B). Induction of IFIT1-luc activity in PMA-treated IFIT1-luc shSAMHD1 THP-1 cells by HIV-1 GFP bearing 75% ΔCA-SP1 was completely absent in STING knock out cells, but maintained in the MAVS knock out cells, consistent with DNA being the predominant viral PAMP detected (Fig. 4A). Confirming these findings, no IRF reporter activity (Fig. 4C) or CXCL-10 production (Fig. 4D) was observed in PMA-treated THP-1 Dual shSAMHD1 cGAS-/- cells infected with HIV-1 75% ΔCA-SP1. Similar findings were also observed for DRV-treated wild type HIV-1 GFP, where induction of IFIT1-luc reporter activity was dependent on STING (Fig. 4E) and cGAS (Fig. 4G), but not MAVS expression (Fig. 4E). Interestingly, whilst CXCL-10 production in these experiments was severely diminished in STING-/-(Fig. 4F) and cGAS-/- (Fig. 4H) cells, levels were also reduced in MAVS-/- cells (Fig. 4F) suggesting a contribution by HIV-1 RNA sensing in the production of this inflammatory cytokine. In all experiments, no significant difference in infection levels between the Ctrl and knockout cell lines was observed (Suppl. Fig. 6A-D).

**Fig. 4.**
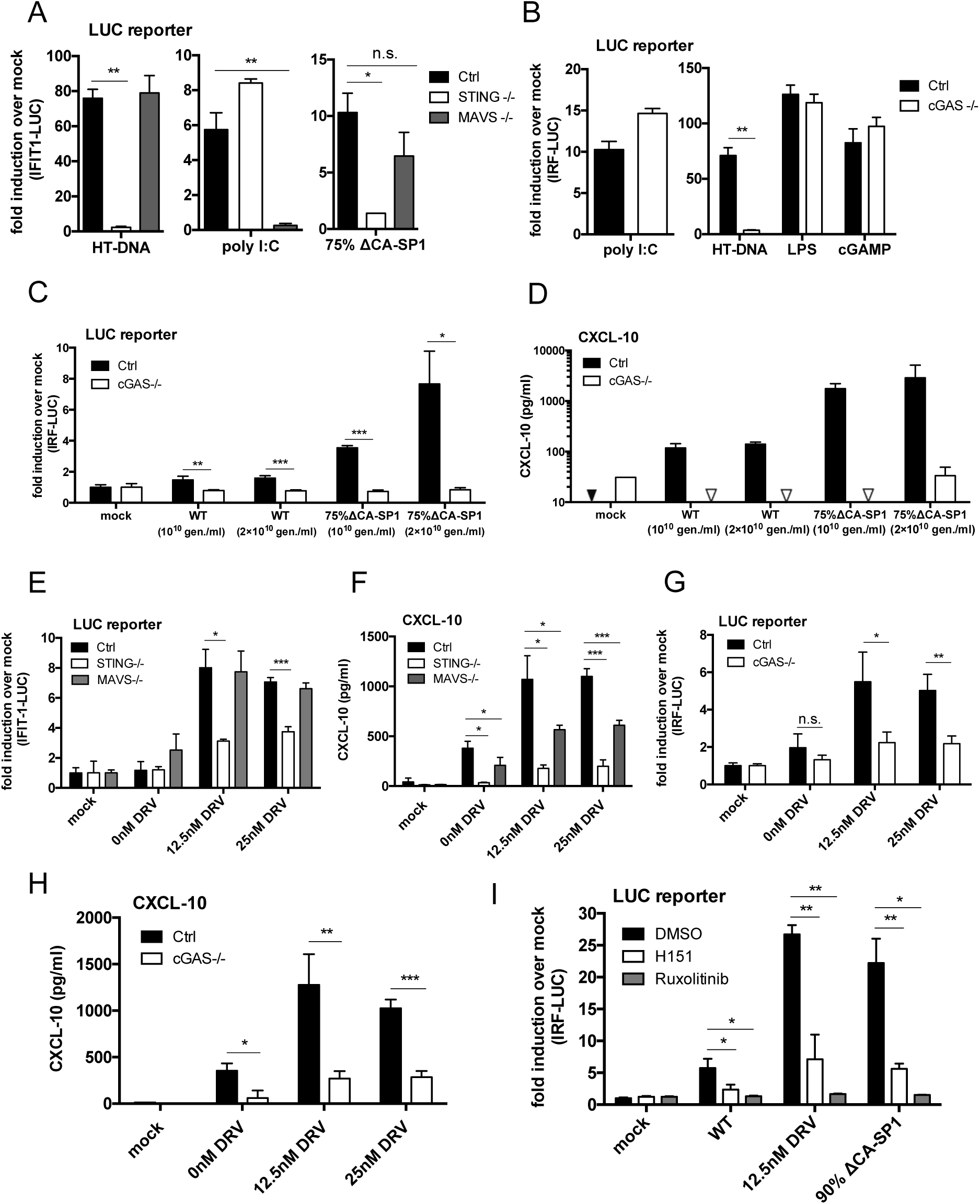
Innate immune activation is DNA sensing-dependent. (A) IFIT-1 reporter activity from PMA-treated (50 ng/ml, 48 h) THP-1-IFIT-1 shSAMHD1 cells lacking STING or MAVS, or a gRNA control (Ctrl) cell line that were transduced for 24 h with HIV-1 GFP 75 % ΔCA-SP1 (0.4 U RT/ml) or stimulated by transfection with either HT-DNA (0.1 µg/ml) or poly I:C (0.5 µg/ml). Gaussia luciferase activity in the supernatant was measured and normalised to mock transduced cells to generate a fold induction. (B-C) IRF reporter activity from PMA-treated (50 ng/ml, 48 h) THP-1 Dual shSAMHD1 cells lacking cGAS or a matching control (Ctrl) cell line that were stimulated for 24 h with poly I:C (transfection, 0.5 µg/ml), HT-DNA (transfection, 0.1 µg/ml), LPS (50 ng/ml) or cGAMP (transfection, 0.5 µg/ml) (B) or transduced for 24 h with increasing doses of HIV-1 GFP containing either 0 % (WT) or 75 % ΔCA-SP1 (1×10^10^ and 2×10^10^ genomes/ml) (C). (D) Level of CXCL-10 protein in the cell supernatants from (C) was measured by ELISA. Triangles indicate CXCL-10 not detected. (E) IFIT-1 reporter activity from PMA-treated (50 ng/ml, 48 h) THP-1-IFIT-1 shSAMHD1 cells lacking STING, MAVS or matching gRNA control (Ctrl) cell line that were transduced for 24 h with DRV-treated HIV-1 GFP as indicated (1×10^10^ genomes/ml). (F) Level of CXCL-10 protein in the cell supernatants from (E) was measured by ELISA. (G) IRF reporter activity from PMA-treated (50 ng/ml, 48 h) THP-1 Dual shSAMHD1 cells lacking cGAS or matching control (Ctrl) cell lines that were transduced for 24 h with DRV-treated HIV-1 GFP as indicated (1×10^10^ genomes/ml). (H) Level of CXCL-10 protein in the cell supernatants from (G) was measured by ELISA. (I) IRF reporter activity from PMA-treated (50 ng/ml, 48 h) THP-1 Dual shSAMHD1 control cells transduced for 48 h with WT, DRV-treated (DRV, 12.5 nM) or HIV-1 GFP containing 90 % ΔCA-SP1 (1×10^10^ genomes/ml) in the presence or absence (DMSO control) of 2 µM ruxolitinib or 0.5 µg/ml H151. Data are presented as mean ± SD of triplicate data repeated 2-4 times. Statistical analyses were performed using the Student’s t-test, with Welch’s correction where appropriate. * *P*<0.05, ** *P*<0.01, *** *P*<0.001. See also Suppl. Fig. 6.

To corroborate data obtained in the CRISPR cell lines, infection assays were also repeated in THP-1 Dual reporter cells in the presence of the recently available STING inhibitor H151 (Haag, Gulen et al., 2018). ISG induction by 12.5 nM DRV-treated or HIV-1 GFP bearing 90% ΔCA-SP1 was greatly reduced by the presence of H151 (Fig. 4I), further supporting a role for DNA sensing in the detection of maturation defective HIV-1. As expected, IRF reporter activity was also suppressed by ruxolitinib (Fig. 4I). Neither H151 nor ruxolitinib affected infection levels in these experiments (Suppl. Fig. 6E).

### Maturation defective viruses fail to saturate TRIM5α in an abrogation-of-restriction assay

If maturation defective viruses consist of defective particles that have a reduced ability to protect viral DNA from cGAS, we hypothesised that these particles may also have a reduced capacity to bind the restriction factor TRIM5α. Rhesus monkey TRIM5α binds HIV-1 capsid and forms hexameric cage-like structures around the intact HIV capsid lattice (Ganser-Pornillos, Chandrasekaran et al., 2011, Li, Chandrasekaran et al., 2016). TRIM5α binding to capsid leads to proteasome recruitment, disassembly of the virus and activation of an innate response (Fletcher, Christensen et al., 2015, Fletcher, Vaysburd et al., 2018, Pertel, Hausmann et al., 2011). Viral restriction can be overcome by co-infection with high doses of a saturating virus in an abrogation-of-restriction assay and this has been suggested to be dependent on the stability of the incoming viral capsid (Jacques, McEwan et al., 2016, Shi & Aiken, 2006).

As a measure of HIV-1 core integrity we tested the ability of the maturation defective viruses to saturate restriction by rhesus macaque TRIM5α. Rhesus FRhK cells were co-infected with a fixed dose of HIV-1 GFP and increasing doses of either wild type untreated HIV-1 luc, LPV-treated HIV-1 luc or HIV-1 luc bearing ΔCA-SP1. Rescue of HIV-1 GFP infectivity from TRIM5α was assessed by flow cytometry measuring GFP positive cells. Viruses that induced a strong innate response, i.e. virus bearing 75 % ΔCA-SP1 mutant (Fig. 5A, Suppl. Fig. 7A) or wild type HIV-1 treated with 30 or 100 nM LPV (Fig. 5B, Suppl. Fig. 7B) showed a reduced ability, or failed to saturate TRIM5α restriction. These data are consistent with cleavage defective HIV-1 particles failing to form the authentic hexameric lattice required for recruitment of TRIM5α (Ganser-Pornillos & Pornillos, 2019, Li et al., 2016) and protection of genome.

**Fig. 5.**
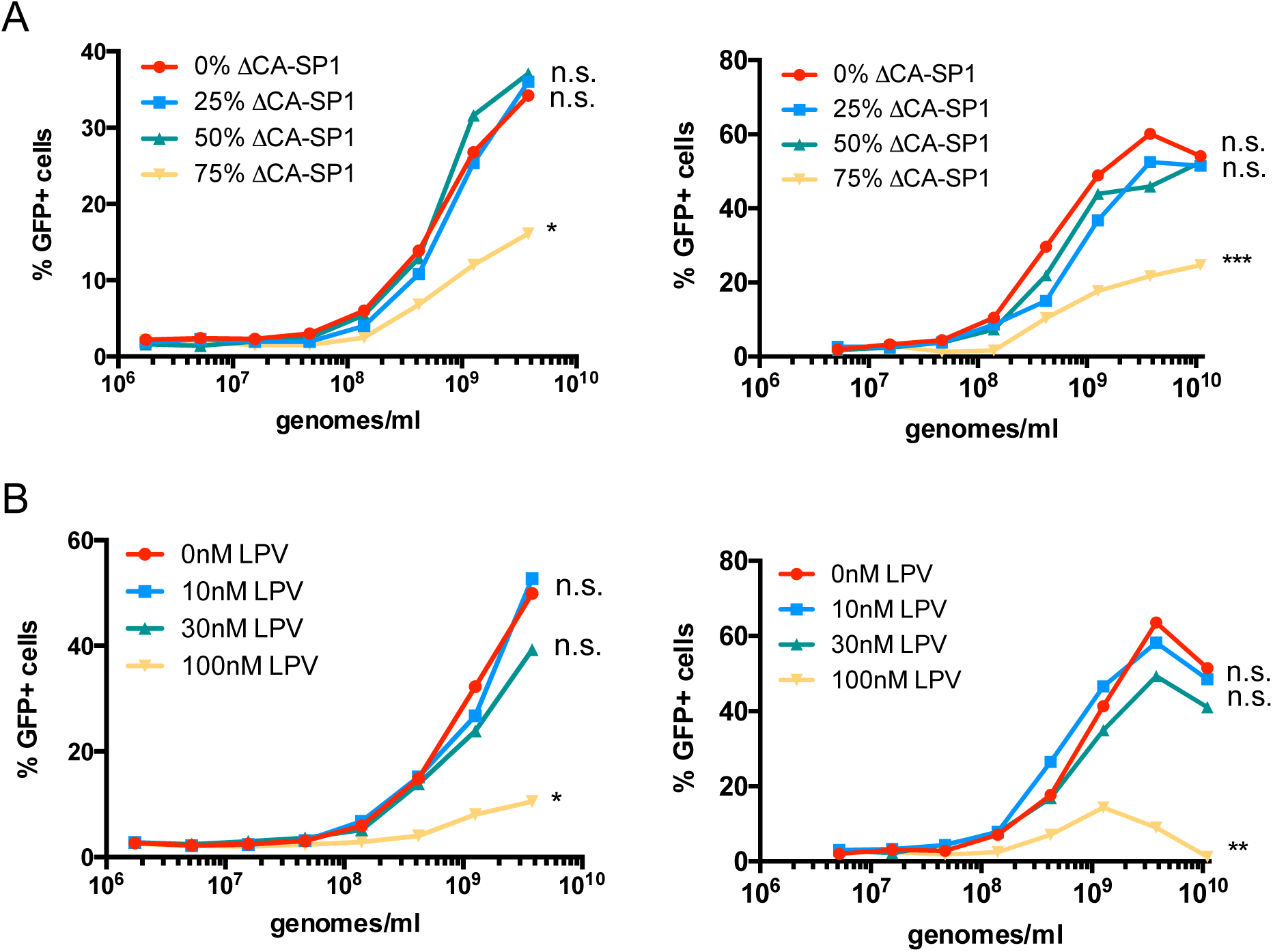
Gag-defective HIV-1 particles are less able to saturate restriction factor TRIM5. (A-B) Abrogation-of-restriction assay in FRhK cells expressing restrictive rhesus TRIM5. FRhK cells were co-transduced with a fixed dose of HIV-1 GFP (5×10^7^ genomes/ml) and increasing doses of HIV-LUC ΔCA-SP1 mutants (A) or LPV-treated HIV-LUC viruses (B) as indicated (1.7×10^6^ - 3.8×10^9^ genomes/ml). Rescue of GFP infectivity was assessed by flow cytometry. Data are presented as singlet % GFP values and two repeats of the experiment are shown. See also Suppl. Fig. 7. Statistical analyses were performed using 2-way ANOVA with multiple comparisons. * *P*<0.05, ** *P*<0.01, *** *P*<0.001.

### Treatment with capsid binding small molecule PF-74 induces HIV-1 to trigger a DNA-sensing dependent ISG response

Recent single molecule analysis of capsid uncoating demonstrated that the capsid binding small molecule inhibitor of HIV, PF-74, accelerates capsid opening (Marquez et al., 2018). We therefore hypothesised that PF-74 treated HIV-1 may activate a DNA-sensing dependent innate immune response. To test this, we infected THP-1 IFIT-1 reporter cells with increasing doses of HIV-1 GFP (0.1-3 U/ml RT) in the presence or absence of 10 µM PF-74. This dose was sufficient to inhibit infection up to 1 U/ml RT HIV-1 GFP, indicating PF-74 is an effective inhibitor of HIV-1, although its potency could be improved (Fig. 6B). Consistent with our hypothesis, at high dose HIV-1 infection (3 U/ml RT) luciferase reporter induction was observed in the presence of PF-74 but not in the DMSO control (Fig. 6A). ISG induction in the presence of 10 µM PF-74 was further confirmed in a second experiment by measuring endogenous *CXCL-10* (Fig. 6C) and *MxA* (Fig. 6D) mRNA expression by qPCR and secreted CXCL-10 by ELISA in the IFIT1-luc reporter cells (Fig. 6E). PF-74 treatment of HIV-1 GFP was further shown to induce a type I IFN response in these cells as IFIT1-luc reporter activity was diminished in the presence of ruxolitinib (Fig. 6F). As expected there was partial inhibition of infection with PF-74 and no difference in infection levels in the presence of ruxolitinib (Suppl. Fig. 8A). Finally, we were able to demonstrate that innate sensing of PF-74-treated HIV-1 was dependent on cGAS as luciferase secretion by PF-74 treated HIV-1 GFP was lost in cGAS-/- cells (Fig. 6G), but maintained in MAVS-/- cells (Fig. 6H). As previously observed, the loss of cGAS (Suppl. Fig. 8B) or MAVS (Suppl. Fig. 8C) had no impact on HIV-1 infectivity suggesting sensing does not contribute to the inhibitory effect of PF74 in these single round infections.

**Fig. 6.**
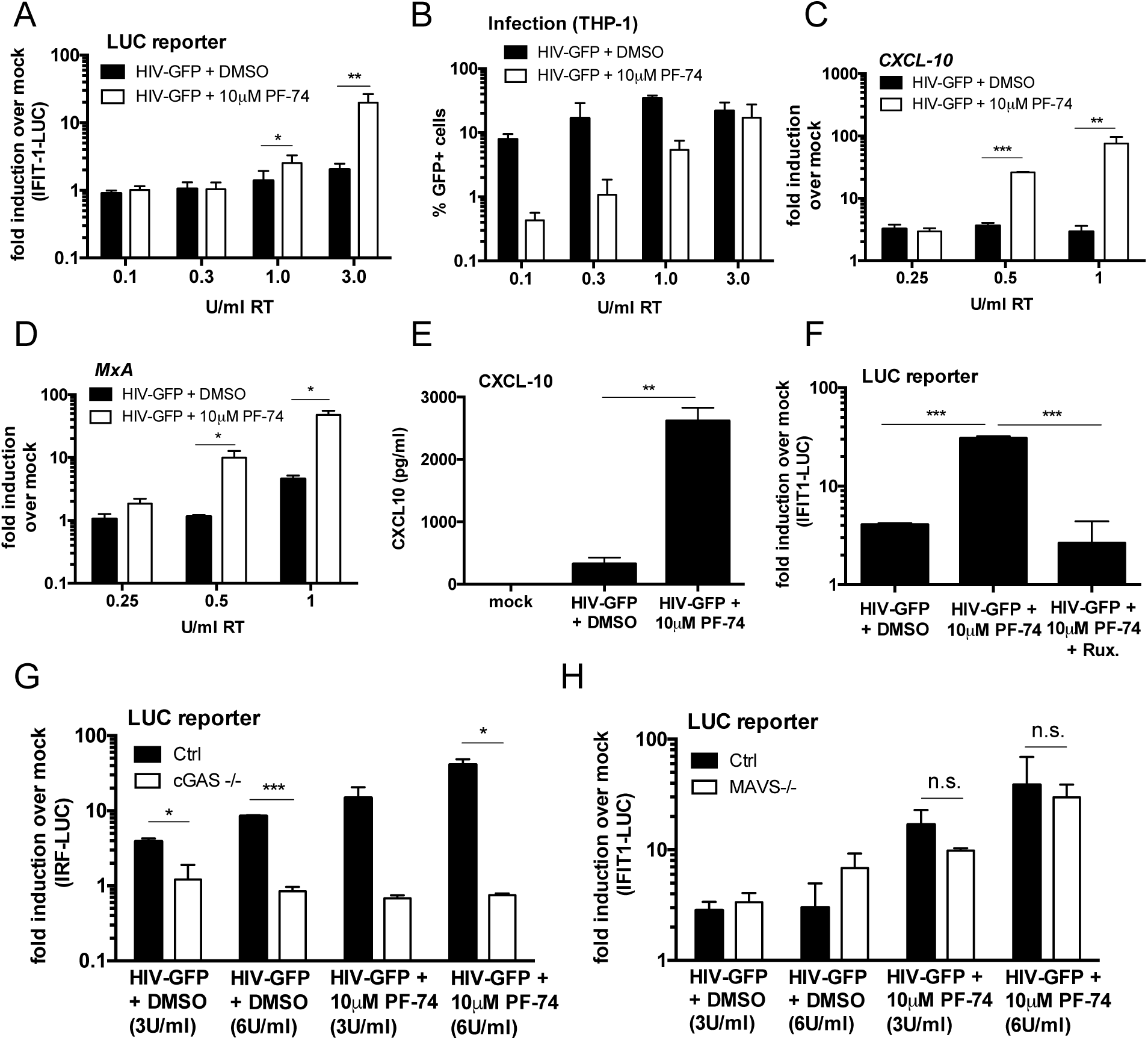
PF-74 treatment induces HIV-1 to trigger a DNA-sensing dependent ISG response. (A) IFIT-1 reporter activity from THP-1-IFIT-1 cells transduced for 24 h with increasing doses (0.1-3 U/ml RT) of HIV-1 GFP in the presence or absence (DMSO control) of PF-74 (10 µM). Gaussia luciferase activity in the supernatant was measured and normalised to mock transduced cells to generate a fold induction. (B) Infection levels of cells from 6A. Cells were analysed for GFP positivity by flow cytometry. (C-D) ISG qPCR from THP-1-IFIT-1 cells transduced for 24 h with increasing doses of HIV-1 GFP (0.25-1 U/ml RT) in the presence or absence (DMSO control) of PF-74 (10 µM). Expression of *CXCL-10* (C) and *MxA* (D) was normalised to an internal control (*GAPDH*) and these values were then normalised to those for the non-transduced mock cells, yielding the fold induction over mock. (E) Level of CXCL-10 protein in the cell supernatants of THP-1-IFIT-1 cells transduced for 24 h with HIV-1 GFP (3 U/ml) in the presence or absence (DMSO control) of PF-74 (10 µM) was measured by ELISA. (F) IFIT-1 reporter activity from THP-1-IFIT-1 cells transduced for 24 h with with HIV-1 GFP (3 U/ml RT) in the presence or absence (DMSO control) of PF-74 (10 µM) and ruxolitinib (Rux, 2 µM) as indicated. (G) IRF reporter activity from THP-1 Dual shSAMHD1 cells lacking cGAS or a matching control (Ctrl) cell line that were transduced for 24 h with increasing doses of HIV-1 GFP (3 U/ml and 6 U/ml) in the presence or absence (DMSO control) of PF-74 (10 µM). (H) IFIT-1 reporter activity from THP-1-IFIT-1 cells lacking MAVS or a matching gRNA control (Ctrl) cell line that were transduced for 24 h with increasing doses of HIV-1 GFP (3 U/ml and 6 U/ml) in the presence or absence (DMSO control) of PF-74 (10 µM). Data are presented as mean ± SD of replicate data (2-6 replicates per condition) repeated at least three times. Statistical analyses were performed using the Student’s t-test, with Welch’s correction where appropriate. * *P*<0.05, ** *P*<0.01, *** *P*<0.001. See also Suppl. Fig. 8.

## Discussion

Effective evasion of innate immune responses is expected to be crucial for successful infection and all viruses have evolved countermeasures to hide PAMPs and/or directly reduce activation of the IFN response (Schulz & Mossman, 2016). Given the small coding capacity of HIV-1 and the general lack of innate activation observed with this virus *in vitro* (Cingoz & Goff, 2019, Lahaye et al., 2013, Rasaiyaah et al., 2013) we had hypothesised that HIV-1 uses its capsid to physically protect nucleic acid PAMPs from innate sensors such as cGAS. In this study we used three approaches to demonstrate that the HIV-1 capsid plays a protective role in preventing IFN induction by viral DNA. By treating HIV-1 with PIs LPV (Fig. 1) or DRV (Suppl. Fig. 3), or mutating the cleavage site between CA and SP1 (Fig. 2) we were able to generate aberrant particles by interfering with capsid maturation. In all cases the resulting viruses had perturbations in Gag cleavage, reduced infectivity (Fig. 1, Fig. 2, Suppl. Fig. 3) and had reduced capacity to saturate the restriction factor TRIM5α in an abrogation-of-restriction assay, indicative of altered stability/capsid integrity (Fig. 5). Importantly, when these viruses were used to infect macrophages they induced a potent IFN response that was not observed on infection with untreated or WT HIV-1 (Fig. 1, Fig. 2, Suppl. Fig. 3). Innate immune responses were almost entirely dependent on viral reverse transcription (Fig. 3) and the cellular DNA sensing machinery comprising cGAS and STING (Fig. 4), consistent with viral DNA being the most important PAMP in these experiments. As a third approach our results were corroborated using the capsid targeting small molecule inhibitor PF-74, which has been proposed to accelerate capsid opening (Marquez et al., 2018). Treatment of HIV-1 with PF-74 also caused a DNA-sensing dependent IFN response (Fig. 6). Together these data support a model in which the WT HIV-1 core remains intact as it traverses the cytoplasm, thus protecting viral DNA from detection by cGAS. Conversely, disruption of capsid maturation or integrity, either chemically or genetically, yields particles that fail to conceal viral DNA and thus activate a cGAS-dependent type 1 IFN response (Suppl. Fig. 9).

We hypothesise that HIV-1 has evolved to cloak viral DNA synthesis within an intact capsid(Jacques et al., 2016, Rasaiyaah et al., 2013). However, a series of studies have reported innate immune activation by WT HIV-1 in macrophages or dendritic cells, but these have required suppression of SAMHD1 by co-transduction with Vpx-containing VLPs(Gao et al., 2013, Johnson, Lucas et al., 2018, Manel, Hogstad et al., 2010, Yoh et al., 2015), high doses of virus(Gao et al., 2013), unpurified viral stocks(Manel et al., 2010, Yan et al., 2010) or manipulation of nuclease TREX1(Yan et al., 2010). A technical complication in testing whether HIV-1, or any other virus, triggers innate immune sensing is controlling for viral dose effects. We and others have found that at very high dose, HIV-1 activates innate immune pathways and this is influenced particularly by whether the viral supernatant is purified. In the experiments presented here, all viruses were DNase treated and purified by centrifugation through sucrose and experiments were designed to control dose between variables. For example, viral dose was normalised by measuring RT activity (SG-PERT) or the number of viral genomes (qPCR) in viral preparations to account for differences in virus production, see legends. Critically, mutating the CA-SP1 cleavage site does not impact RT activity and treatment with protease inhibitors only inhibited supernatant RT activity at the highest dose used, and only by a few fold. We propose that small ISG responses to the doses of WT HIV-1 used here are likely due to low frequency uncoating events because the minimal ISG response to WT HIV-1 was also dependent on cGAS (Fig. 4C, Fig. 6G).

Further support for the important role of capsid in innate immune evasion comes from data in dendritic cells demonstrating that unlike wild type HIV-1, wild type HIV-2 activates a strong RT- and cGAS-dependent IFN response. This difference in innate activation mapped to capsid (Lahaye et al., 2013). Why the HIV-2 capsid, unlike the capsid of HIV-1, fails to protect RT products from innate sensors is the subject of ongoing investigation, but given that HIV-2 does not replicate in dendritic cells and macrophages (Chauveau, Puigdomenech et al., 2015, Duvall, Lore et al., 2007) these observations suggest that evasion of sensing by cGAS is a necessary requirement for replication in myeloid cells.

Interestingly, we discovered here that single round infection triggering of IFN did not lead to reduction in viral infectivity. This was particularly apparent in experiments using the JAK1/2 inhibitor ruxolitinib, which potently reduced the ISG response to viruses with defective capsids, but had no impact on GFP positivity of the cells (Suppl. Fig. 4C, Suppl. Fig. 6E, Suppl. Fig. 8A). Similarly, cGAS knockout severely blunted ISG responses, but did not lead to a corresponding increase in GFP positivity (Suppl. Fig. 6B, D, Suppl. Fig. 8B). We propose that during single round infections the virus has already integrated by the time IFN is produced and GFP expression is not particularly sensitive to its anti-viral effects. Indeed, the IFITM proteins (OhAinle, Helms et al., 2018, Petrillo, Thorne et al., 2018, Yu & Liu, 2018), TRIM5α (OhAinle et al., 2018, Pertel et al., 2011) and MxB (Goujon, Moncorge et al., 2013, Kane, Yadav et al., 2013, OhAinle et al., 2018) are the major IFN-induced inhibitors of HIV-1 in THP-1 cells and are not expected to impact GFP expression.

Another interesting finding that warrants further investigation is the observation that MAVS contributed to CXCL-10 production in response to infection with DRV-treated virus (Fig. 4F), but did not contribute to the corresponding IFIT-1 reporter activity (Fig. 4E). MAVS-dependent pathways are known to activate transcription factors other than IRF-3, such as NF-κB (Seth, Sun et al., 2005), which also contributes to the production of CXCL-10 (Yeruva, Ramadori et al., 2008), but not activation of the IFIT-1 reporter (Grandvaux, Servant et al., 2002). It is therefore possible that activation of MAVS by HIV-1 contributes to NF-κB activation in these cells but not an IRF-3 response.

In summary these findings highlight the crucial role of the HIV-1 capsid in masking viral nucleic acids from innate immune sensors, particularly in protecting viral DNA from detection by cGAS/STING. As such, disrupting capsid integrity through mutation, treatment with protease inhibitors, or the capsid targeting small molecule PF-74 yields viral particles that fail to shield their PAMPs and thus activate a potent IFN response that is not observed with the WT virus. Together these data suggest that the therapeutic activity of capsid or protease - targeting therapeutics, for example the recently described HIV-1 capsid inhibitor from Gilead Sciences(Yant, Mulato et al., 2019), may be enhanced by induction of local antiviral IFN responses *in vivo* that could contribute to viral clearance by the innate and adaptive immune system. Furthermore these findings encourage the design of therapeutics targeting capsids or structural proteins generally, which may also benefit from unmasking viral PAMPs and induction of innate immune responses.

## Acknowledgments

We thank Veit Hornung for kindly providing THP-1-IFIT-1 cells. This work was funded through a Wellcome Trust Senior Biomedical Research Fellowship (GJT), the European Research Council under the European Union’s Seventh Framework Programme (FP7/2007-2013)/ERC (grant HIVInnate 339223) and the National Institute for Health Research University College London Hospitals Biomedical Research Centre and a Wellcome Trust Collaborative award.

## Author Contributions

RPS and GJT conceived the study. RPS, LH, TP, ET, MS and LZA performed the experiments. RPS, LH, TP and GJT analysed the data. RPS and GJT wrote the manuscript.

## Declaration of Interests

The authors declare no competing interests.

## Methods

### Cells and reagents

HEK293T and U87 cells were maintained in DMEM (Gibco) supplemented with 10 % foetal bovine serum (FBS, Labtech) and 100 U/ml penicillin plus 100 µg/ml streptomycin (Pen/Strep; Gibco). THP-1 cells were maintained in RPMI (Gibco) supplemented with 10 % FBS and Pen/Strep. THP-1-IFIT-1 cells that had been modified to express Gaussia luciferase under the control of the *IFIT-1* promoter were described previously (Mankan et al., 2014). THP-1 Dual Control and cGAS-/- cells were obtained from Invivogen. Lopinavir (LPV), darunavir (DRV), nevirapine (NVP), raltegravir, zidovudine (AZT) and tenofovir (TDF) were obtained from AIDS reagents. STING inhibitor H151 was obtained from Invivogen. JAK inhibitor ruxolitinib was obtained from CELL guidance systems. PF-74 was obtained from Sigma. Lipopolysaccharide, IFNβ and poly I:C were obtained from Peprotech. Sendai virus was obtained from Charles River Laboratories. Herring-testis DNA was obtained from Sigma. cGAMP was obtained from Invivogen. For stimulation of cells by transfection, transfection mixes were prepared using lipofectamine 2000 according to the manufacturer’s instructions (Invitrogen).

### Generation of ΔCA-SP1, RT D185E and INT D116N viruses

pLAIΔEnv GFP/Luc ΔCA-SP1 (Gag mutant L363I M367I) was generated by two rounds of site-directed mutagenesis (using Pfu Turbo DNA polymerase, Agilent) using primers: LAI_Gag_L363I fwd: 5’ CCGGCCATAAGGCAAGAGTTATCGCTGAAGCAATG 3’

LAI_Gag_L363I rev: 5’ GTTACTTGGCTCATTGCTTCAGCGATAACTCTTGC 3’

LAI_Gag_M367I fwd: 5’ GCAAGAGTTATCGCTGAAGCAATCAGCCAAGTAAC 3’

LAI_Gag_M367I rev: 5’ GTAGCTGAATTTGTTACTTGGCTGATTGCTTCAGC 3’

pLAIΔEnv GFP and pLAIΔEnv GFP ΔCA-SP1 RT D185E and INT D116N were generated by site-directed mutagenesis using the following primers:

LAI_ RT D185E fwd: 5’ ATAGTTATCTATCAATACATGGAAGATTTGTATG 3’

LAI_ RT D185E rev: 5’ AAGTCAGATCCTACATACAAATCTTCCATGTATTG 3’

LAI_ INT D116N fwd: 5’ GGCCAGTAAAAACAATACATACAAACAATGGCAGC 3’

LAI_ INT D116N rev: 5’ ACTGGTGAAATTGCTGCCATTGTTTGTATGTATTG 3’

In all cases mutated sequences were confirmed by sequencing, excised by restriction digestion and cloned back into the original plasmid.

### Isolation of primary monocyte-derived macrophages

Primary monocyte-derived macrophages (MDM) were prepared from fresh blood from healthy volunteers. The study was approved by the joint University College London/University College London Hospitals NHS Trust Human Research Ethics Committee and written informed consent was obtained from all participants. Peripheral blood mononuclear cells (PBMCs) were isolated by density gradient centrifugation using Lymphoprep (Stemcell Technologies). PBMCs were washed three times with PBS and plated to select for adherent cells. Non-adherent cells were washed away after 1.5 h and the remaining cells incubated in RPMI (Gibco) supplemented with 10 % heat-inactivated pooled human serum (Sigma) and 40 ng/ml macrophage colony stimulating factor (R&D systems). Cells were further washed after 3 days and the medium changed to RPMI supplemented with 10 % heat-inactivated FBS. MDM were then infected 3-4 days later. Replicate experiments were performed with cells derived from different donors.

### Editing of cells by CRISPR/Cas 9

THP-1 IFIT-1 shSAMHD1 STING-/- and MAVS-/- cells were previously described (Tie et al., 2018). Briefly, lentiparticles to generate CRISPR/Cas9-edited cell lines were produced by transfecting 10 cm dishes of HEK293T cells with 1.5 µg of plentiCRISPRv2 encoding gene specific guide RNAs (Addgene plasmid #52961), 1 µg of p8.91 packaging plasmid (Zufferey, Nagy et al., 1997), and 1 µg of vesicular stomatitis virus-G glycoprotein expressing plasmid pMDG (Genscript) using Fugene 6 transfection reagent (Promega) according to the manufacturer’s instructions. Virus supernatants were harvested at 48 and 72 h post-transfection, pooled and used to transduce THP-1 IFIT-1 shSAMHD1 cells by spinoculation (1000 *xg*, 1 h, room temperature). Transduced cells were selected using puromycin (1 µg/ml, Merck Millipore) and single clones isolated by limiting dilution in 96 well plates. Clones were screened for successful gene knock out by luciferase assay and immunoblotting. gRNA sequences:

STING: TCCATCCATCCCGTGTCCCAGGG

MAVS: CAGGGAACCGGGACACCCTC

Non-targeting control: ACGGAGGCTAAGCGTCGCAA

### Production of virus in 293T cells

HIV-1 and lentiviral particles were produced by transfection of HEK293T cells in T150 flasks using Fugene 6 transfection reagent (Promega) according to the manufacturer’s instructions. For full length HIV-1 with a BaL envelope cells were transfected with 8.75 µg pR9.BaL per flask. For HIV-1 GFP/Luc each flask was transfected with 2.5 µg of vesicular stomatitis virus-G glycoprotein expressing plasmid pMDG (Genscript) and 6.25 µg pLAIΔEnv GFP/Luc. Virus supernatants were harvested at 48 and 72 h post-transfection, pooled, DNase treated (2 h at 37 °C, DNaseI, Sigma) and subjected to ultracentrifugation over a 20 % sucrose cushion. Viral particles were finally resuspended in RPMI supplemented with 10 % FBS. For production of viruses in the presence of lopinavir or darunavir, the inhibitors were added at 24 h post-transfection and replaced after harvest at 48 h. Lentiparticles for SAMHD1 depletion were generated as previously described (Georgana, Sumner et al., 2018). Viruses were titrated by infecting U87 cells (10^5^ cells/ml) or PMA-treated THP-1 cells (2×10^5^ cells/ml) with dilutions of sucrose purified virus in the presence of polybrene (8 µg/ml, Sigma) for 48 h and enumerating GFP-positive cells by flow cytometry using the FACS Calibur (BD) and analysing with FlowJo software.

### SG-PERT

Reverse transcriptase activity of virus preparations was quantified by qPCR using a SYBR Green-based product-enhanced RT (SG-PERT) assay as described (Vermeire, Naessens et al., 2012).

### Genome copy/RT products measurements

For viral genome copy measurements RNA was extracted from 2 µl sucrose purified virus using the RNeasy mini kit (QIAgen). The RNA was then treated with TURBO DNase (Thermo Fisher Scientific) and subjected to reverse transcription using Superscript III reverse transcriptase and random hexamers according to the manufacturer’s protocol (Invitrogen). Genome copies were then measured by Taqman qPCR using primers against GFP (see below). For RT product measurements DNA was extracted from 5×10^5^ infected cells using the DNeasy Blood & Tissue kit (QIAgen) according to the manufacturer’s protocol. DNA concentration was quantified using a Nanodrop for normalisation. RT products were quantified by Taqman qPCR using TaqMan Gene Expression Master Mix (ThermoFisher) and primers and probe specific to GFP. A dilution series of plasmid encoding GFP was measured in parallel to generate a standard curve to calculate the number of GFP copies.

*GFP* fwd: 5’- CAACAGCCACAACGTCTATATCAT -3’

*GFP* rev: 5’- ATGTTGTGGCGGATCTTGAAG -3’

*GFP* probe: 5’- FAM-CCGACAAGCAGAAGAACGGCATCAA-TAMRA -3’

### Infection assays

THP-1 cells were infected at a density of 2×10^5^ cells/ml. For differentiation THP-1 cells were treated with 50 ng/ml phorbol 12-myristate 13-acetate (PMA, Peprotech) for 48 h. Luciferase reporter assays were performed in 24 well plates and qPCR and ELISA in 12 well plates. Infection levels were assessed at 48 h post-infection through enumeration of GFP positive cells by flow cytometry. Infections in THP-1 cells were performed in the presence of polybrene (8 µg/ml, Sigma). Input dose of virus was normalised either by RT activity (measured by SG-PERT) or genome copies (measured by qPCR) as indicated.

### Luciferase reporter assays

Gaussia/Lucia luciferase activity was measured by transferring 10 µl supernatant to a white 96 well assay plate, injecting 50 µl per well of coelenterazine substrate (Nanolight Technologies, 2 µg/ml) and analysing luminescence on a FLUOstar OPTIMA luminometer (Promega). Data were normalised to a mock-treated control to generate a fold induction.

### ISG qPCR

RNA was extracted from THP-1/primary MDM using a total RNA purification kit (Norgen) according to the manufacturer’s protocol. Five hundred ng RNA was used to synthesise cDNA using Superscript III reverse transcriptase (Invitrogen), also according to the manufacturer’s protocol. cDNA was diluted 1:5 in water and 2 µl was used as a template for real-time PCR using SYBR® Green PCR master mix (Applied Biosystems) and a Quant Studio 5 real-time PCR machine (Applied Biosystems). Expression of each gene was normalised to an internal control (*GAPDH*) and these values were then normalised to mock-treated control cells to yield a fold induction. The following primers were used:

*GAPDH:* Fwd 5’-GGGAAACTGTGGCGTGAT-3’, Rev 5’-GGAGGAGTGGGTGTCGCTGTT-3’ *CXCL-10:* Fwd 5’-TGGCATTCAAGGAGTACCTC-3’, Rev 5’-TTGTAGCAATGATCTCAACACG-3’ *IFIT-2:* Fwd 5’-CAGCTGAGAATTGCACTGCAA-3’, Rev 5’-CGTAGGCTGCTCTCCAAGGA-3’ *MxA:* Fwd 5’-ATCCTGGGATTTTGGGGCTT-3’, Rev 5’-CCGCTTGTCGCTGGTGTCG-3’ *CXCL-2:* Fwd 5’-GGGCAGAAAGCTTGTCTCAA-3’, Rev 5’-GCTTCCTCCTTCCTTCTGGT-3’

### ELISA

Cell supernatants were harvested for ELISA at 24 h post-infection/stimulation and stored at - 80 °C. CXCL-10 protein was measured using Duoset ELISA reagents (R&D Biosystems) according to the manufacturer’s instructions.

### Immunoblotting

For immunoblotting of viral particles 2×10^11^ genome copies of virus were boiled for 10 min in 6X Laemmli buffer (50 mM Tris-HCl (pH 6.8), 2 % (w/v) SDS, 10% (v/v) glycerol, 0.1% (w/v) bromophenol blue, 100 mM β-mercaptoethanol) before separating on 4-12 % Bis-Tris polyacrylamide gradient gel (Invitrogen). For immunoblot analysis of THP-1 cells, 3×10^6^ cells were lysed in a cell lysis buffer containing 50 mM Tris pH 8, 150 mM NaCl, 1 mM EDTA, 10% (v/v) glycerol, 1 % (v/v) Triton X100, 0.05 % (v/v) NP40 supplemented with protease inhibitors (Roche), clarified by centrifugation at 14,000 x *g* for 10 min and boiled in 6X Laemmli buffer for 5 min. Proteins were separated by SDS-PAGE on 12 % polyacrylamide gels. After PAGE, proteins were transferred to a Hybond ECL membrane (Amersham biosciences) using a semi-dry transfer system (Biorad). Primary antibodies were from the following sources: mouse anti-β-actin (Abcam), rabbit-anti-SAMHD1 (Proteintech) and mouse-anti-HIV-1capsid p24 (183-H12-5C, AIDS Reagents). Primary antibodies were detected with goat-anti-mouse/rabbit IRdye 800CW infrared dye secondary antibodies and membranes imaged using an Odyssey Infrared Imager (LI-COR Biosciences).

### Abrogation of restriction assay

FRhK cells were plated in 48 well plates at 5×10^4^ cells/ml. The following day cells were co-transduced in the presence of polybrene (8 µg/ml, Sigma) with a fixed dose of HIV-1 GFP (5×10^7^ genome copies/ml) and increasing doses of HIV-LUC ΔCA-SP1 mutants or LPV-treated HIV-LUC viruses (1.7×10^6^ - 3.8×10^9^ genome copies/ml). Rescue of GFP infectivity was assessed 48 h later by flow cytometry using the FACS Calibur (BD) and analysing with FlowJo software.

### Statistical analyses

Statistical analyses were performed using an unpaired Student’s t-test (with Welch’s correction where variances were unequal) or a 2-way ANOVA with multiple comparisons, as indicated. * *P*<0.05, ** *P*<0.01, *** *P*<0.001.

---

**Suppl. Fig. 1.**
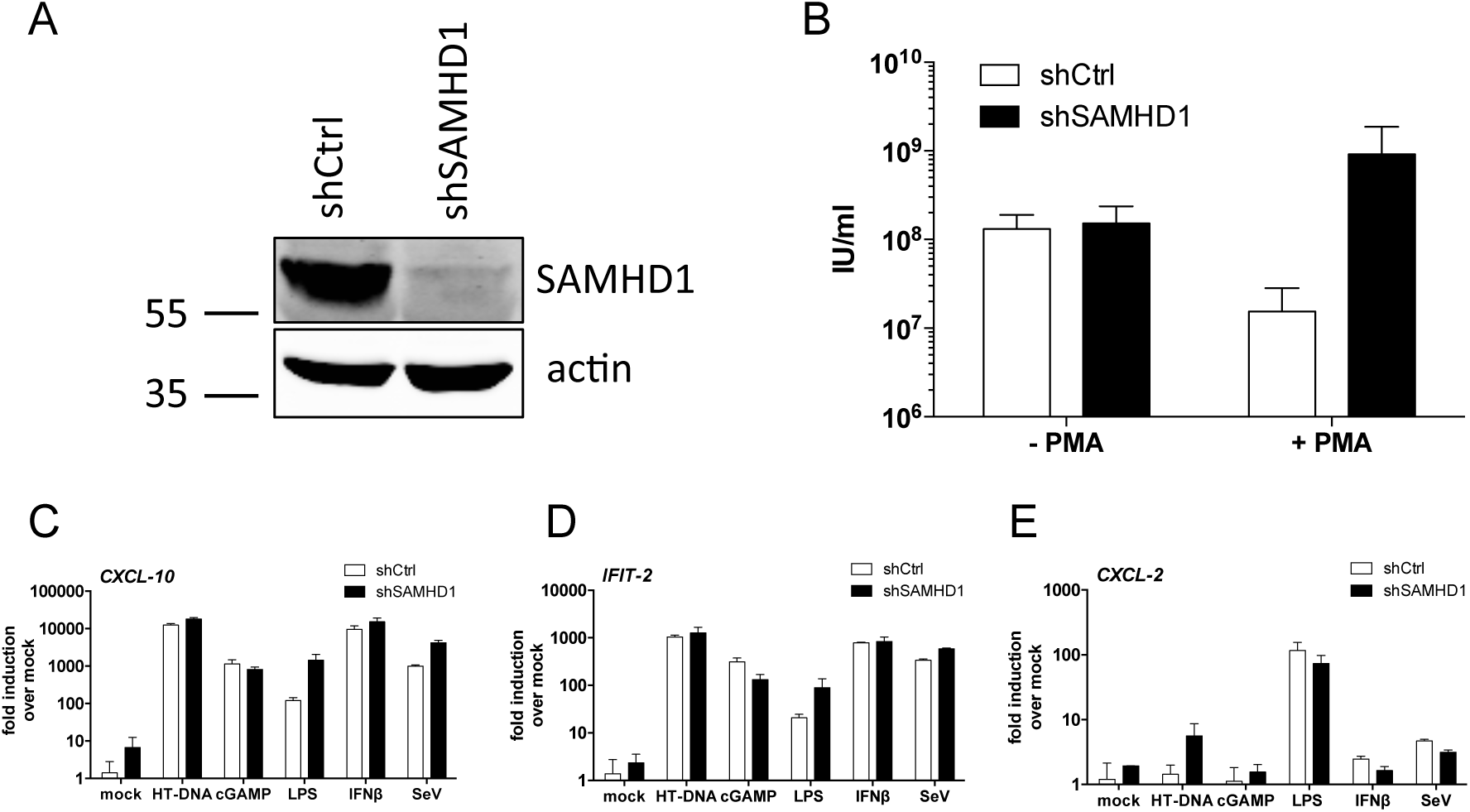
Generation of THP-1 cell lines depleted for SAMHD1. (A) Immunoblot of THP-1 cell line with stable depletion of SAMHD1 (shSAMHD1) or a corresponding control shRNA (shCtrl) (see Methods). Blots were probed with anti-SAMHD1 and anti-actin antibodies. Molecular mass markers (in kDa) are indicated on the left. (B) Titration of HIV-1 GFP on THP-1 shCtrl and THP-1 shSAMHD1 cells that had been treated or not with PMA (50 ng/ml) for 48 h. Infectious units per ml (IU/ml) were calculated by enumeration of GFP-positive cells by flow cytometry. Data are presented as mean ± SD from three separate titrations performed in singlet. (C-E) ISG qPCR from PMA-treated (50 ng/ml, 48 h) THP-1 shCtrl and THP-1 shSAMHD1 cells that had been stimulated for 8 h with HT-DNA (20 ng/ml, transfected), cGAMP (1 µg/ml), LPS (1 µg/ml), IFNβ (10 ng/ml) or Sendai virus (SeV) (0.2 HA U/ml). Cells were lysed for RNA extraction, cDNA was synthesised and then used for qPCR analysis. Expression of *CXCL-10* (C), *IFIT-2* (D) and *CXCL-2* (E) was normalised to an internal control (*GAPDH*) and these values were then normalised to those for the non-stimulated mock cells, yielding the fold induction over mock. Data are presented as mean ± SD of duplicate data repeated at least twice.

**Suppl. Fig. 2.**
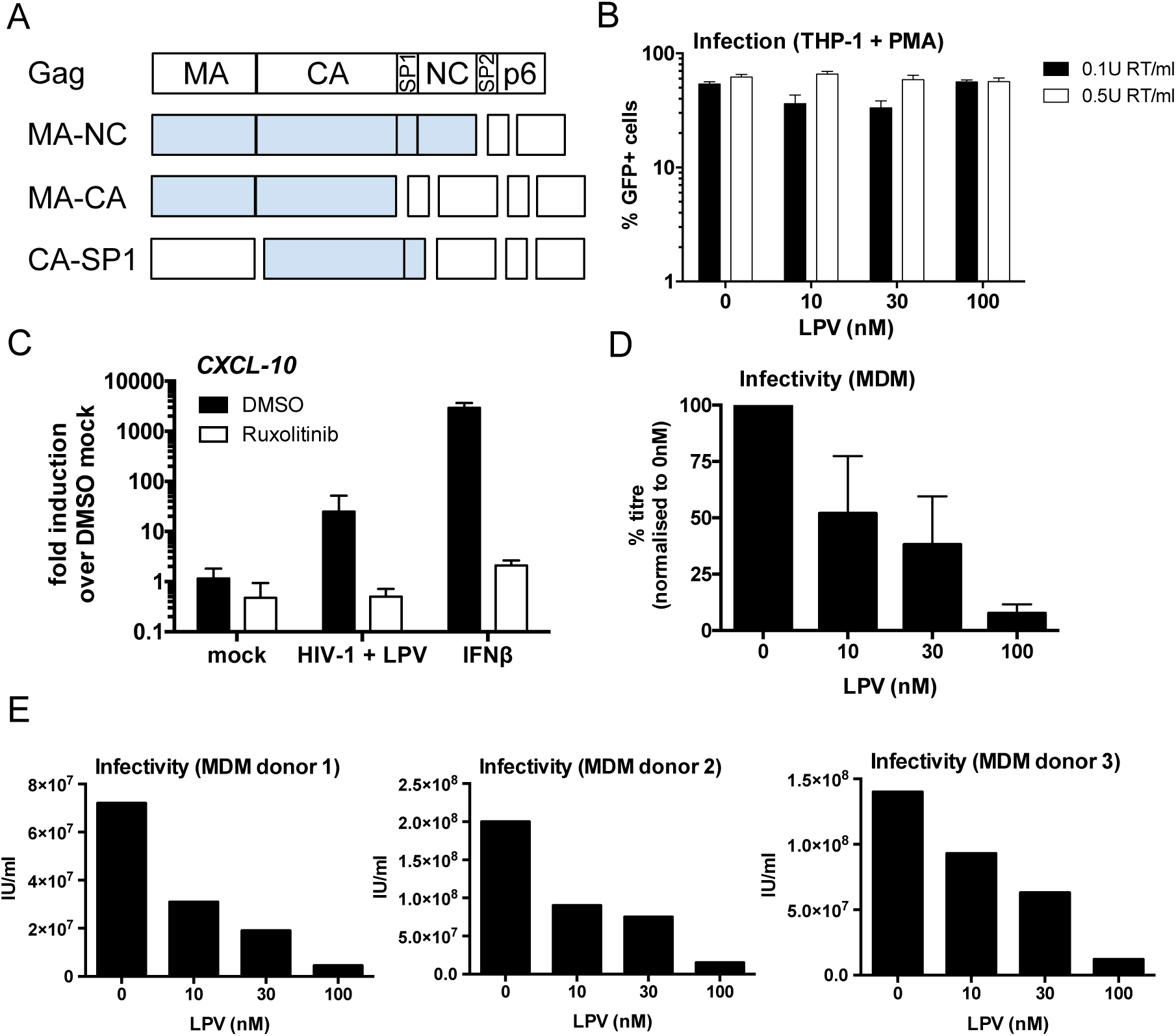
PI treatment induces HIV-1 to trigger an ISG response in macrophages. (A) Schematic of intermediate Gag cleavage products. MA: matrix, CA: capsid, SP1: spacer peptide 1, NC: nucleocapsid, SP2: spacer peptide 2. (B) Infection levels of cells from Fig. 1D-G. PMA-treated (50 ng/ml, 48 h) THP-1 shSAMHD1 cells were transduced for 48 h with LPV-treated HIV-1 GFP viruses (0.1 U RT/ml or 0.5 U RT/ml). Cells were analysed for GFP positivity by flow cytometry. Data are presented as mean ± SD of triplicate data repeated at least three times. (C) ISG qPCR from PMA-treated (50 ng/ml, 48 h) THP-1 shSAMHD1 cells that had been transduced for 24 h with 0.2 U RT/ml HIV-1 GFP that had been treated with 30 nM LPV, or stimulated with 1 ng/ml IFNβ as a control, in the presence or absence (DMSO control) or 2 µM ruxolitinib. Expression of *CXCL-10* was normalised to an internal control (*GAPDH*) and these values were then normalised to those for the non-transduced, DMSO-treated mock cells, yielding the fold induction over DMSO mock. Data are presented as mean ± SD of triplicate data repeated at least twice. (D-E) Titration of LPV-treated R9 BaL viruses on primary monocyte-derived macrophages. Infectious units per ml (IU/ml) were calculated by staining cells with a FITC-labelled anti-p24 antibody and analysing by flow cytometry. Data from individual donors are represented in E and collated in D, represented as percentage titre normalised to R9 BaL produced in the absence of LPV (0 nM). Data are shown as mean ± SD.

**Suppl. Fig. 3.**
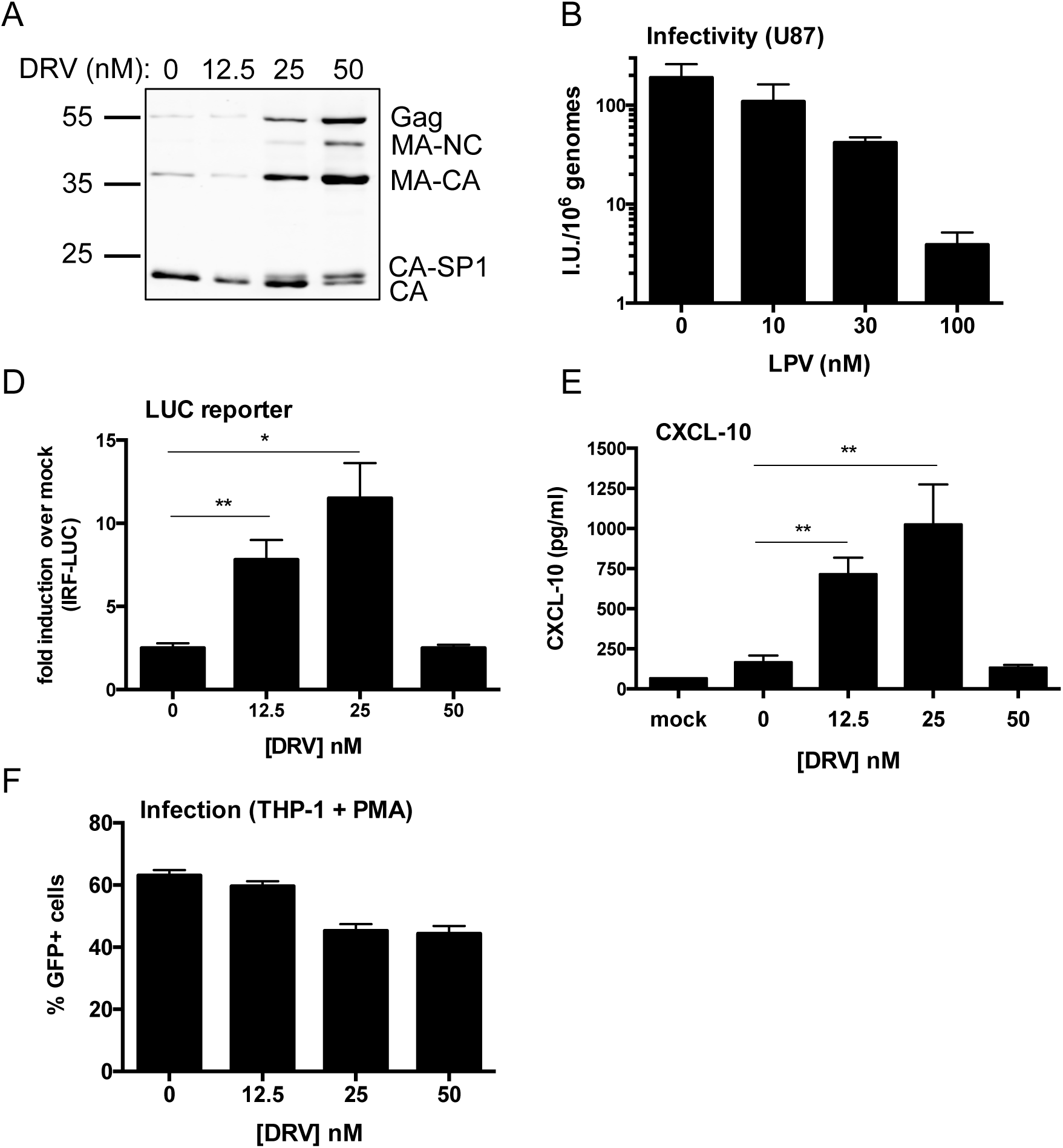
Treatment of HIV-1 with darunavir also triggers an ISG response. (A) Immunoblot of HIV-1 GFP virus particles (2×10^11^ genomes) produced in HEK293T cells in the presence of increasing doses of protease inhibitor darunavir (DRV, 0-50 nM) and then purified through a 20 % sucrose cushion. The blot was probed with an anti-HIV-1 p24 antibody. Molecular mass markers (in kDa) are indicated on the left and Gag cleavage products on the right. (B) Titration of DRV-treated HIV-1 GFP viruses on U87 cells. Infectious units per ml (IU/ml) were calculated by enumeration of GFP-positive cells by flow cytometry. These data were then normalised to genomes/ml data calculated by qPCR to account for variations in viral production. Data are presented as mean ± SD from three separate titrations performed in singlet. (D) IRF reporter activity from PMA-treated (50 ng/ml, 48 h) THP-1 Dual shSAMHD1 cells transduced for 24 h with DRV-treated HIV-1 GFP (1×10^10^ genomes/ml). Gaussia luciferase activity in the supernatant was measured and normalised to mock transduced cells, yielding a fold induction. (E) Level of CXCL-10 protein in the cell supernatant from D was measured by ELISA. (F) Infection levels of cells from D-E. Cells were analysed for GFP positivity by flow cytometry at 48 h post-transduction. Data for D-F are presented as mean ± SD of triplicate data repeated at least twice. Statistical analyses were performed using the Student’s t-test, with Welch’s correction where appropriate and comparing to the 0 nM DRV condition. * *P*<0.05, ** *P*<0.01.

**Suppl. Fig. 4.**
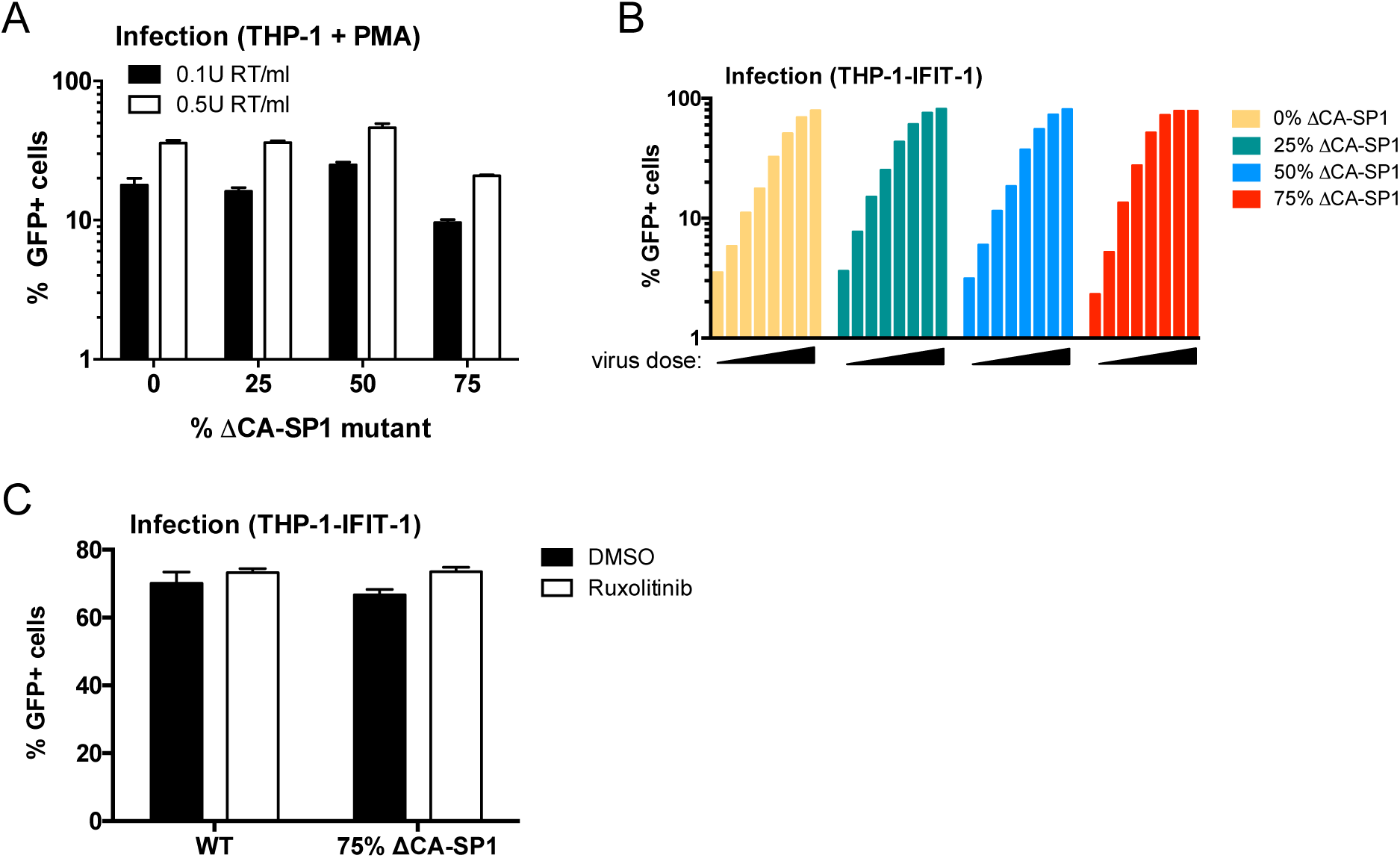
HIV-1 with Gag protease cleavage mutation induces ISGs in THP-1 cells. (A) Infection levels of cells from Fig. 2D-F. PMA-treated (50 ng/ml, 48 h) THP-1 shSAMHD1 cells were transduced for 48 h with HIV-1 GFP ΔCA-SP1 viruses (0.1 U RT/ml or 0.5 U RT/ml). Cells were analysed for GFP positivity by flow cytometry. (B) Infection levels of cells from Fig. 2G. THP-1-IFIT-1 cells were transduced for 48 h with HIV-1 GFP ΔCA-SP1 viruses (0.016 – 0.2 U RT/ml). Cells were analysed for GFP positivity by flow cytometry. (C) Infection levels of cells from Fig. 2H. THP-1-IFIT-1 cells were transduced for 48 h with HIV-1 GFP containing either 0 % (WT) or 75 % ΔCA-SP1 mutant in the presence or absence (DMSO control) of 2 µM ruxolitinib. Cells were analysed for GFP positivity by flow cytometry. Data are shown as individual measurements (B) or mean ± SD from triplicate data (A, C) repeated at least three times.

**Suppl. Fig. 5.**
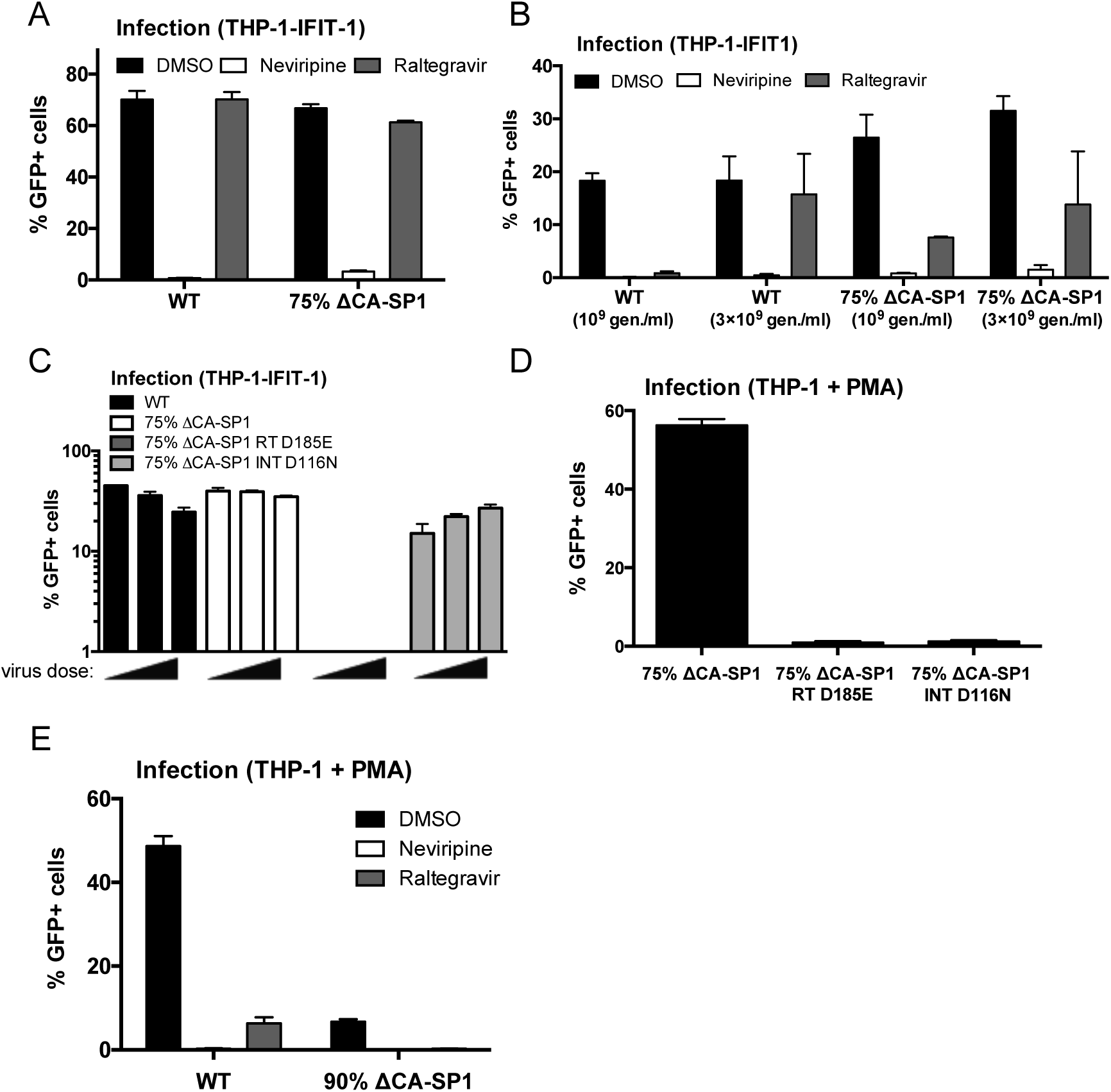
Innate immune activation is RT-dependent. (A) Infection levels from Figures 3A and B. THP-1-IFIT-1 cells were transduced for 48 h with HIV-1 GFP containing 0 % (WT) or 75 % ΔCA-SP1 mutant (1 U RT/ml) in the presence or absence (DMSO control) of 5 µM neviripine or 10 µM raltegravir. Cells were analysed for GFP positivity by flow cytometry. (B) Infection levels for Figures 3C and D. THP-1-IFIT-1 cells were transduced for 48 h with increasing doses of 0 % (WT) or 75 % ΔCA-SP1 mutant (10^9^ ^and^ 3×10^9^ genomes/ml) in the presence or absence (DMSO control) of 5 µM neviripine or 10 µM raltegravir. Cells were analysed for GFP positivity by flow cytometry. (C) Infection levels for Figures 3E and F. THP-1-IFIT-1 cells were transduced for 48 h with with increasing doses of HIV-1 GFP containing 0 % ΔCA-SP1 (WT), 75 % ΔCA-SP1, 75 % ΔCA-SP1 carrying a mutation in reverse transcriptase (75 % ΔCA-SP1 RT D185E) or 75 % ΔCA-SP1 carrying a mutation in integrase (75 % ΔCA-SP1 INT D116N) (3.75×10^9^, 7.5×10^9^ and 1.5×10^10^ genomes/ml). Cells were analysed for GFP positivity by flow cytometry. (D) PMA-treated (50 ng/ml, 48 h) THP-1 Dual shSAMHD1 cells were transduced for 48 h with 75 % ΔCA-SP1, 75 % ΔCA-SP1 carrying a mutation in reverse transcriptase (75 % ΔCA-SP1 RT D185E) or 75 % ΔCA-SP1 carrying a mutation in integrase (75 % ΔCA-SP1 INT D116N) (3×10^9^ genomes/ml). Cells were analysed for GFP positivity by flow cytometry. (E) PMA-treated (50 ng/ml, 48 h) THP-1 Dual shSAMHD1 control cells were transduced for 48 h with WT HIV-1 GFP or 90 % ΔCA-SP1 mutant (1×10^10^ genomes/ml) in the presence or absence (DMSO control) of 5 µM neviripine or 10 µM raltegravir. Data are presented as mean ± SD of triplicate data repeated 2-3 times.

**Suppl. Fig. 6.**
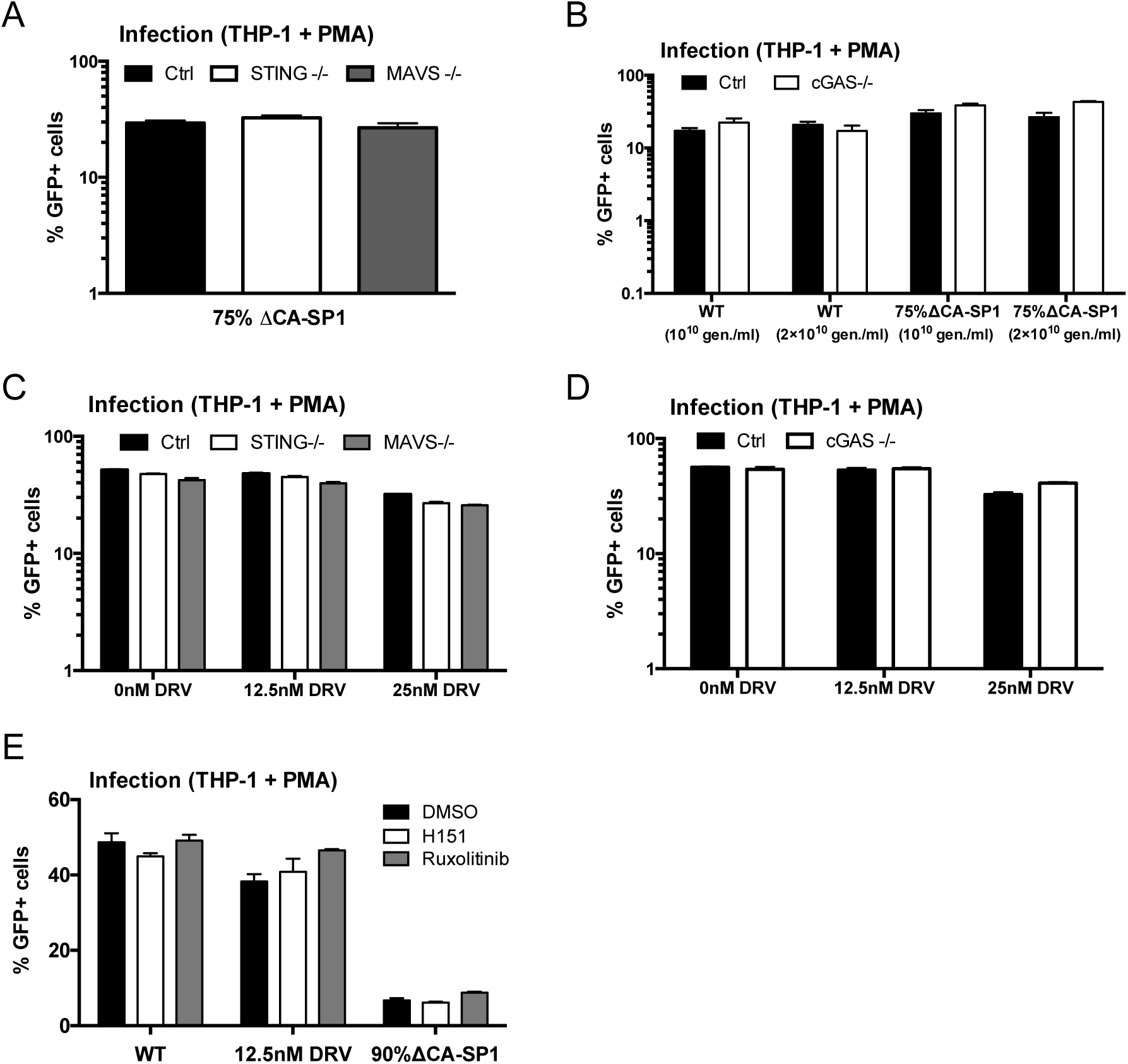
Innate immune activation is DNA sensing-dependent. (A) Infection levels of cells from Fig. 4A. PMA-treated (50 ng/ml, 48 h) THP-1-IFIT-1 shSAMHD1 cells lacking STING or MAVS, or a gRNA control (Ctrl) cell line that were transduced for 48 h with HIV-1 GFP 75 % ΔCA-SP1 (0.4 U RT/ml). (B) Infection levels of cells from Fig. 4 C-D. PMA-treated (50 ng/ml, 48 h) THP-1 Dual shSAMHD1 cells lacking cGAS or a matching control (Ctrl) cell line were transduced for 48 h with increasing doses of HIV-1 GFP containing either 0 % (WT) or 75 % ΔCA-SP1 (1×10^10^ and 2×10^10^ genomes/ml). (C) Infection levels of cells from Fig. 4E-F. PMA-treated (50 ng/ml, 48 h) THP-1-IFIT-1 shSAMHD1 cells lacking STING, MAVS or matching gRNA control (Ctrl) cell lines were transduced for 48 h with DRV-treated HIV-1 GFP as indicated (1×10^10^ genomes/ml). (D) Infection levels of cells from Fig. 4G-H. PMA-treated (50 ng/ml, 48 h) THP-1 Dual shSAMHD1 cells lacking cGAS or matching control (Ctrl) cell lines were transduced for 48 h with DRV-treated HIV-1 GFP as indicated (1×10^10^ genomes/ml). (E) Infection levels of cells from Fig. 4I. PMA-treated (50 ng/ml, 48 h) THP-1 Dual shSAMHD1 control cells were transduced for 48 h with WT, DRV-treated (DRV, 12.5 nM) or HIV-1 GFP 90 % ΔCA-SP1 (1×10^10^ genomes/ml) in the presence or absence (DMSO control) of 2 µM ruxolitinib or 0.5 µg/ml H151. Cells were analysed for GFP positivity by flow cytometry. Data are presented as mean ± SD of triplicate data repeated 2-4 times.

**Suppl. Fig 7.**
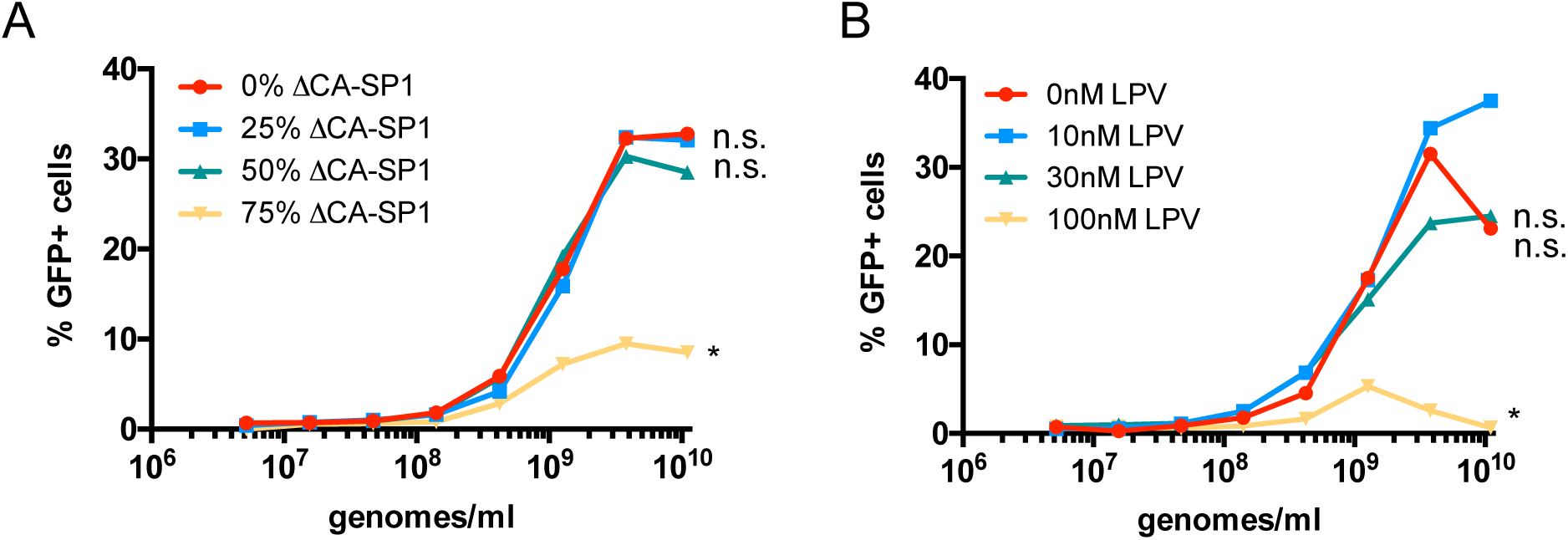
Gag-defective HIV-1 particles are less able to saturate restriction factor TRIM5. Abrogation-of-restriction assay in FRhK cells expressing restrictive rhesus TRIM5. (A) Repeat assay of data presented in Fig. 5A. FRhK cells were co-transduced with a fixed dose of HIV-1 GFP (5×10^7^ genomes/ml) and increasing doses of HIV-LUC ΔCA-SP1 mutants as indicated (5.2×10^6^ - 1.1×10^10^ genomes/ml). (B) Repeat assay of data presented in Fig. 5B. FRhK cells were co-transduced with a fixed dose of HIV-1 GFP (5×10^7^ genomes/ml) and increasing doses of LPV-treated HIV-LUC viruses as indicated (5.2×10^6^ - 1.1×10^10^ genomes/ml). Rescue of GFP infectivity was assessed by flow cytometry. Data are presented as singlet % GFP values. Statistical analyses were performed using 2-way ANOVA with multiple comparisons. * *P*<0.05.

**Suppl. Fig. 8.**
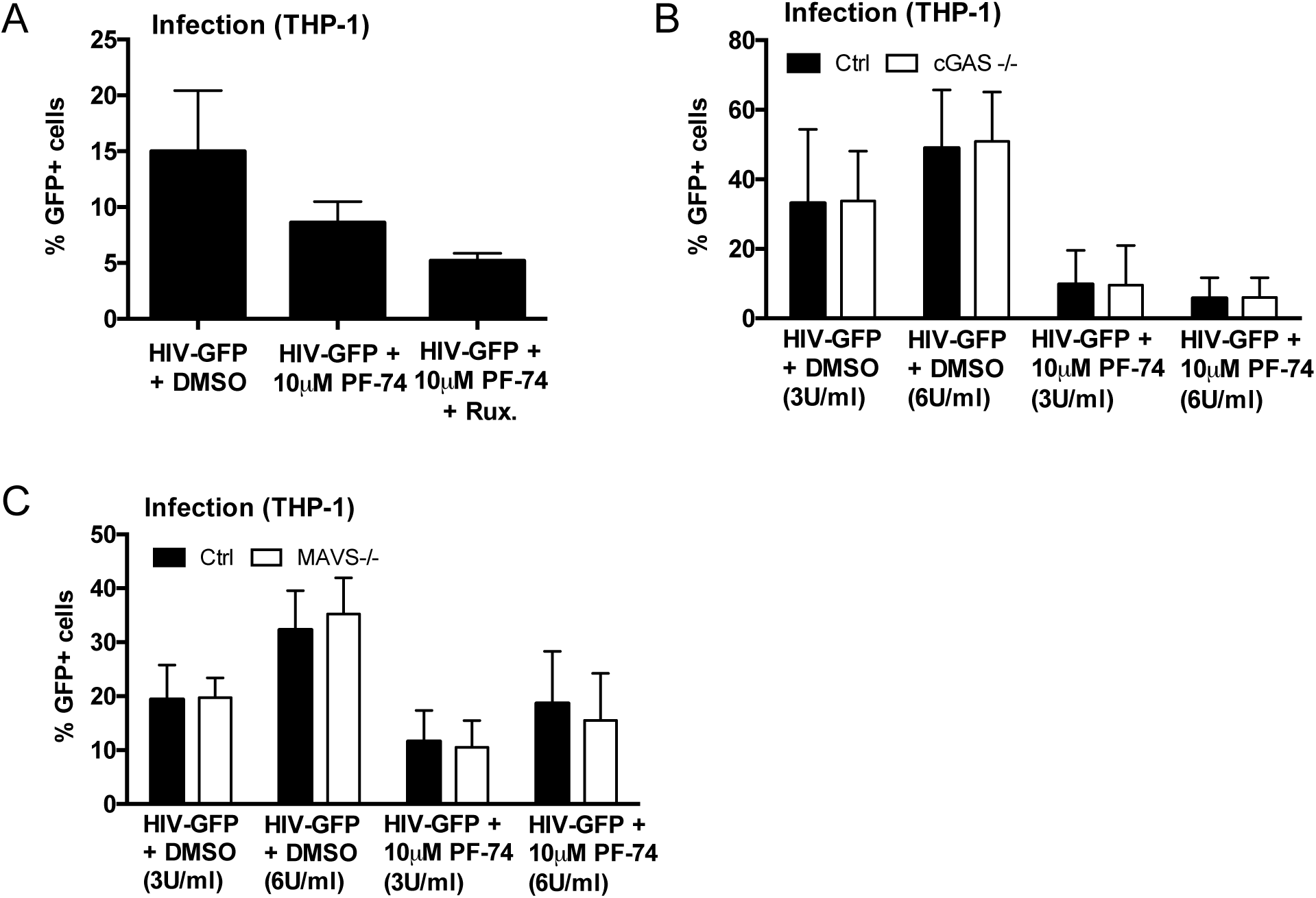
PF-74 treatment induces HIV-1 to trigger a DNA-sensing dependent ISG response. (A) Infection levels of cells from Fig. 6F. THP-1-IFIT-1 cells were transduced for 48 h with with HIV-1 GFP (3 U/ml RT) in the presence or absence (DMSO control) of PF-74 (10 µM) and ruxolitinib (Rux, 2 µM) as indicated. Cells were analysed for GFP positivity by flow cytometry. (B) Infection levels of cells from Fig. 6G. THP-1 Dual shSAMHD1 cells lacking cGAS or a matching control (Ctrl) cell line were transduced for 48 h with increasing doses of HIV-1 GFP (3 U/ml and 6 U/ml) in the presence or absence (DMSO control) of PF-74 (10 µM). Cells were analysed for GFP positivity by flow cytometry. (C) Infection levels of cells from Fig. 6H. THP-1-IFIT-1 cells lacking MAVS or a matching gRNA control (Ctrl) cell line were transduced for 48 h with increasing doses of HIV-1 GFP (3 U/ml and 6 U/ml) in the presence or absence (DMSO control) of PF-74 (10 µM). Cells were analysed for GFP positivity by flow cytometry. Data are presented as mean ± SD of replicate data (2-4 replicates per condition) repeated at least three times.

**Suppl. Fig. 9.**
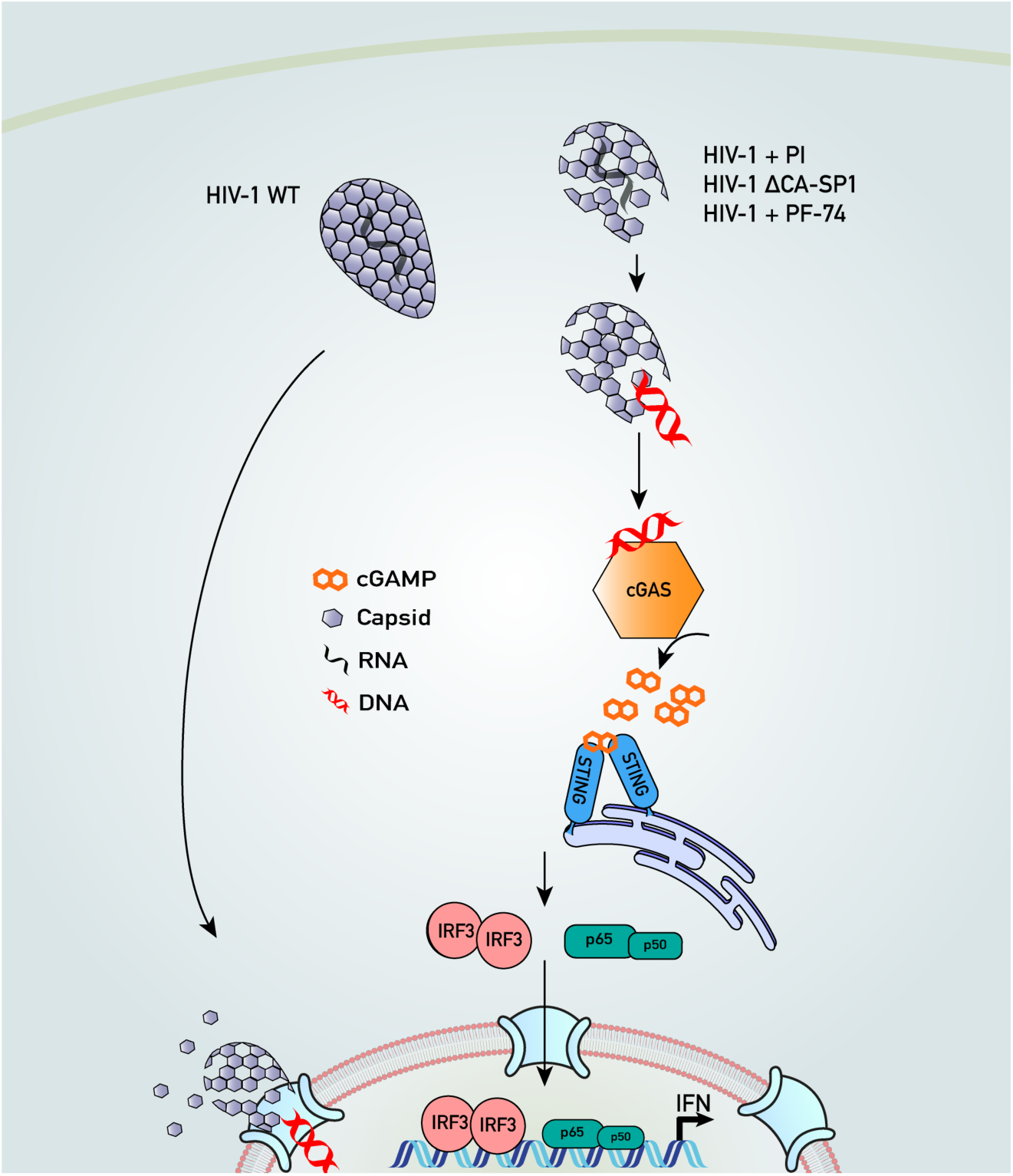
The HIV-1 capsid protects viral DNA from sensing by cGAS. Disrupting HIV-1 capsid formation causes cGAS sensing of viral DNA. After entry wild-type (WT) HIV-1 stays intact as it traverses the cytoplasm allowing it to synthesise its DNA without activating a type I IFN response. Conversely treatment of HIV-1 with protease inhibitors (PI), capsid destabilising small molecule PF-74 or mutation of the protease cleavage site between capsid and spacer peptide 1 (HIV-1 ΔCA-SP1) leads to defective particles that fail to protect viral DNA from innate sensor cGAS. Binding of cGAS to viral DNA leads to the production of cGAMP that binds STING and stimulates IFN production through activation of the transcription factors IRF3 and p50/p65 (NF-κB).

